# A long noncoding RNA, *LOC157273*, is the effector transcript at the chromosome 8p23.1-*PPP1R3B* metabolic traits and type 2 diabetes risk locus

**DOI:** 10.1101/2020.03.24.000620

**Authors:** Alisa K. Manning, Anton Scott Goustin, Erica L. Kleinbrink, Pattaraporn Thepsuwan, Juan Cai, Donghong Ju, Aaron Leong, Miriam S. Udler, James Bentley Brown, Mark O. Goodarzi, Jerome I. Rotter, Robert Sladek, James B. Meigs, Leonard Lipovich

**Affiliations:** Clinical and Translational Epidemiology Unit, Mongan Institute, Massachusetts General Hospital, Boston, MA, USA; Department of Medicine, Harvard Medical School, Boston, MA, USA; Programs in Metabolism and Medical & Population Genetics, Broad Institute of MIT and Harvard, Cambridge, MA, USA; Center for Molecular Medicine & Genetics, Wayne State University, Detroit, MI, USA; Karmanos Cancer Institute at Wayne State University, Detroit, MI, USA; Center for Human Genetics Research, Massachusetts General Hospital, Boston, MA, USA; Division of General Internal Medicine, Massachusetts General Hospital, Boston, MA, USA; Diabetes Unit, Massachusetts General Hospital, Boston, MA, USA; Department of Statistics, University of California, Berkeley, CA, USA; Centre for Computational Biology, University of Birmingham, B15 2TT Birmingham, United Kingdom; Division of Endocrinology, Diabetes, and Metabolism, Department of Medicine, Cedars-Sinai Medical Center, Los Angeles, CA, USA; The Institute for Translational Genomics and Population Sciences, Departments of Pediatrics and Medicine, Los Angeles Biomedical Research Institute at Harbor-UCLA Medical Center, Torrance, CA USA; Department of Human Genetics, McGill University, Montréal, Québec, Canada; Department of Medicine, McGill University, Montréal, Québec, Canada; McGill University and Genome Québec Innovation Centre, Montréal, Québec, Canada; Department of Neurology, School of Medicine, Wayne State University, Detroit, MI, USA

**Keywords:** insulin resistances, hepatic glycogen storage, long non-coding RNA, metabolism, type 2 diabetes, regulatory mechanisms

## Abstract

**Aims:** Causal transcripts at genomic loci associated with type 2 diabetes are mostly unknown. The chr8p23.1 variant rs4841132, associated with an insulin resistant diabetes risk phenotype, lies in the second exon of a long non-coding RNA (lncRNA) gene, *LOC157273*, located 175 kilobases from *PPP1R3B*, which encodes a key protein regulating insulin-mediated hepatic glycogen storage in humans. We hypothesized that *LOC157273* regulates expression of *PPP1R3B* in human hepatocytes.

**Methods:** We tested our hypothesis using Stellaris fluorescent in-situ hybridization to assess subcellular localization of *LOC157273*; siRNA knockdown of *LOC157273*, followed by RT-PCR to quantify *LOC157273* and *PPP1R3B* expression; RNA-seq to quantify the whole-transcriptome gene expression response to *LOC157273* knockdown and an insulin-stimulated assay to measure hepatocyte glycogen deposition before and after knockdown.

**Results:** We found that siRNA knockdown decreased *LOC157273* transcript levels by approximately 80%, increased *PPP1R3B* mRNA levels by 1.7-fold and increased glycogen deposition by >50% in primary human hepatocytes. An A/G heterozygous carrier (vs. three G/G carriers) had reduced *LOC157273* abundance due to reduced transcription of the A allele and increased PPP1R3B expression and glycogen deposition.

**Conclusion:** We show that the lncRNA *LOC157273* is a negative regulator of *PPP1R3B* expression and glycogen deposition in human hepatocytes and the causal transcript at an insulin resistant type 2 diabetes risk locus.

## Introduction

Type 2 diabetes (T2D), a continually growing scourge worldwide, arises from the interaction of multiple factors with genetic susceptibility in insulin sensitivity and secretion pathways to increase risk (1-3). The search for genetic determinants of T2D and its risk factors has revealed over 400 common variants at over 250 coding and regulatory genomic loci that influence multiple distinct aspects of type 2 diabetes pathophysiology (4-8).

In quantitative trait genome-wide association studies (GWAS) of non-diabetic individuals, we showed that the minor (rs4841132-A) allele at the chromosome 8p23.1 variant rs4841132 (NR_040039.1:n.548A>G; reference allele A has frequency ∼11%) was significantly associated with an insulin resistance phenotype characterized by increased levels of fasting glucose (FG) and insulin (FI), elevated levels of triglycerides and an increased waist-hip ratio (4). This chromosome 8 locus is highly pleiotropic; and rs4841132 and nearby SNPs have been consistently associated with increased T2D risk as well as T2D-related metabolic phenotypes including glycemia in pregnancy, obesity, HDL:LDL ratio, total cholesterol, triglycerides, c-reactive protein levels, coronary artery disease, subclinical atherosclerosis, and fatty liver disease (4, 9-15).

The variant rs4841132 resides ∼175 kb from the nearest protein-coding gene, *PPP1R3B. PPP1R3B* encodes the glycogen-targeting subunit of PP1 protein phosphatase and is expressed most strongly in liver in both rodents and man; and at lower levels in skeletal muscle and other tissues (16-18). PPP1R3B connects ambient insulin to hepatic glycogen regulation: its overexpression in hepatocytes markedly increases both basal and insulin-stimulated glycogen synthesis (19). PPP1R3B has long been an attractive target for diabetes therapy, based on the concept of tipping ambient glycemic balance towards hepatic glycogen deposition (17, 20).

We previously localized rs4841132 to exon 2 of a previously unannotated long non-coding RNA (lncRNA) gene, *LOC157273* (ENSG00000254235.1; NR_040039.1) (21). In ancestry-specific analyses we identified a second variant rs9949 (chr8 distance 189084 kb; r^2^_YRI_ with rs4841132, 0.18; r^2^_CEU_, 0.01) that resided in the second exon of *PPP1R3B* and was weakly associated with FI (P=6.9×10^−5^) (21) and T2D (p-value 5.9×10^−4^) in African ancestry individuals. The lncRNA encoded at *LOC157273* is a plausible effector transcript for both variants, as lncRNAs are highly enriched at trait- and disease-associated loci, a few are now known to regulate metabolic pathways and disease risk, and many lncRNAs are *cis*-regulators, exerting both positive and negative regulation of neighboring protein-coding genes (22-28).

These observations support a potential genetic regulatory relationship between *LOC157273* and *PPP1R3B* that could explain the observed metabolic trait and T2D risk GWAS associations at the chr8p23.1 locus (29, 30). As *PPP1R3B* is abundantly expressed in the human liver where it regulates glycogen storage, we studied cultured human hepatocytes to test the hypothesis that *LOC157273* regulates *PPP1R3B* expression, and consequently insulin-mediated glycogen deposition, and that *LOC157273* regulation of *PPP1R3B* varies by genotype at rs4841132.

## Methods

### SNP Genotyping

We obtained primary human hapatocytes from commercial sources and genotyped them by sequencing the region surrounding rs4841132 in a 2.9 kb LOC157273 amplicon from purified DNA (**Supplementary Text)**. We identified one rs4841132 A/G heterozygote out of 16 available hepatocyte donors (**Supplementary Table S1, Supplementary Figure S1)**. This study was not considered human subjects research, as the research was performed using de-identified biospecimans from deceased individuals that were commercially obtained from Lonza (formerly Triangle Research Labs) or ThermoFisher Scientific (formally LifeTech).

### Cellular localization of LOC157273 with Stellaris RNA fluorescent in situ hybridization (FISH)

We used a custom-synthesized 48-probe set (LGC Biosearch Technologies; Petaluma, CA) of non-overlapping fluorescent-tagged oligonucleotides that tiled the 3.4 kb LOC157273 transcript. Probe nucleotide choices at all polymorphic sites were based on the NR_040039.1 reference transcript, which contains the rs4841132-A (minor) allele. Fixed cells grown on collagen-coated glass coverslips were probed with the pooled probe set following the Biosearch Technologies Stellaris FISH protocol for adherent cells (https://www.biosearchtech.com/support/resources/stellaris-protocols). After mounting using Vectashield with DAPI, the hybridized coverslips were examined under an AxioObserver inverted fluorescence microscope (Carl Zeiss Microscopy) equipped with a 63×/1.40 oil objective lens. Red bodies in the merged images denote the Quasar 570 signal from the LOC157273 molecules; the blue-colorized DAPI staining shows cell nuclei. Greater detail is provided in the Supplement.

### TaqMan quantitative reverse-transcriptase PCR (qRT-PCR) to measure the expression levels of two isoforms of PPP1R3B mRNA and a single isoform of LOC157273 lncRNA

Oligo(dT) priming was used for reverse transcription of RNA into cDNA for all Taqman qRT-PCR analyses. Primers and probe-sets are described in more detail in the Supplement and Supplementary Table S2. The *LOC157273* amplicon spans intron 1 and includes 36 nt of exon 1 and 41 nt of exon 2. *PPP1R3B* transcription was assessed with two probe sets—one for the hepatocyte-specific mRNA, and one for the (more ubiquitous) mRNA. Three or four biological replicates were obtained for Taqman qRT-PCR in primary human hepatocyte donors TRL4079, Hu8200, TRL4056B, TRL4105A and TRL4108. All reactions were run as technical triplicates using the Applied Biosystems 7500 Fast Real-Time Instrument.

### Small interfering (si) RNA knockdown of LOC157273

Small-interfering (si)RNA knockdown of *LOC157273* was performed on primary human hepatocytes from donor Hu8200 (genotype G/G at rs4841132) plated into collagen-coated 6-well plates (ThermoFisher Scientific #A11428-01). Cells were incubated in complete Williams’ E medium and transfected with siRNA (50 nM; Dharmacon) and Lipofectamine® RNAiMAX reagent (ThermoFisher Scientific #13778075) in a final concentration of 1 mL/well Opti-MEM™. The target sequences in the 3.4 kb *LOC157273* lncRNA (NR_040039.1 or Ensembl ENST00000520390.1) were: siRNA09 in the 3’-end of the third exon (GGGAAGGGTTAGAGAGGTC), siRNA11 in the first exon (CAACTTAGCTTCTCCATTTTT), siRNA13 near the 5’-end of the third exon (AGAGAAGGACTGAAGATCATT) and siRNA15 in the second exon (TCAGAGGACTTGACACCAT) where the sequences represent the sense-strand DNA targeted by the siRNAs. Six hours after transfection, 1 mL of complete medium (including 1X HepExtend) was pipetted into each well. On the next day, medium was removed after gentle up-and-down-pipetting (tritutation) to dislodge dead cells and the transfected monolayer was supplemented first with 1 mL of complete Williams’ E and then with ice-cold complete medium (including HepExtend) containing 1 mg/mL fibronectin (ThermoFisher Scientific Geltrex #A1413202), followed by gentle trituration to mix the fibronectin (final concentration becomes 0.5 mg/mL). Complete medium with HepExtend was changed daily in the evening (with gentle trituration) until 120 hr after initial plating. We performed pooled analysis of TaqMan qRTPCR results from 3 biological replicates of the siRNA knockdown experiment after applying sequential corrections, including log transformation, mean centering, and autoscaling (54).

### Transcriptome-wide effects of LOC157273 knockdown using RNA sequencing

Transcriptome sequencing was performed by the Broad Institute’s Sequencing Platform (31). Using the NCBI refGen database (hg19) and R/Bioconductor packages (GenomicFeatures, rsubread), we collapsed transcript annotations into genes and obtained gene counts using the pairedEnd option in featureCounts for all experimental treatment groups. Three biologic replicates were performed using primary human hepatocytes (Hu8200 donor) for 4 different siRNAs (siRNA09, siRNA11, siRNA13, siRNA15) and for control experiments consisting of either mock or scrambled siRNA transfection. Differentially expressed genes were obtained with DESeq2, with adjustment for batch effects and normalization for small gene counts in a model comparing gene counts in siRNA11 and siRNA15 experimental conditions to mock and scramble controls. We observed low counts which is standard when analyzing transcriptome-wide expression of protein-coding genes and lncRNAs even in the absence of siRNA knockdown. Therefore, we utilized shrinkage based on re-estimating the variance using dispersion estimates with a negative binomial distribution (32-34). We performed hierarchical clustering with normalized gene counts (after removing batch effects) and Reactome pathway-based analysis (both restricted to genes with P<0.001 and effects as large as the *PPP1R3B* effect), and gene-set enrichment analysis (for all genes) with ReactomePA (35).

### Glycogen Deposition Assay in response to insulin or glucagon

We developed a protocol to measure glycogen content in cultured primary human hepatocytes using donor TRL4079 (heterozygote A/G at rs4841132) and donors TRL4055A, TRL4113 and TRL4012 (homozygous G/G at rs4841132.) In the assay, adapted from Gómez-Lechón et al. (36), Aspergillus niger amyloglucosidase (Sigma-Aldrich #A7420) degrades cell-derived glycogen to glucose, which was measured in a fluorescent peroxide/peroxidase assay. Cells were lysed with ice-cold solubilization buffer (2% CHAPS, 150 mM NaCl, 25 mM Tris-HCl, pH 7.2) containing 1X HALT™ protease inhibitor cocktail (ThermoFisher 78430), using 400 μL of lysis buffer for each well of a 6-well plate. To measure glucose polymerized as glycogen, the lysate was diluted 1:10 with sodium acetate buffer (50 mM Na-acetate, pH 5.5) and treated with either 0.75-1.5 U amyloglucosidase (Sigma A7420) at pH 5.5 (60 min at 37°C) or pH 5.5 buffer without enzyme. After incubation, 5 μL aliquots of the ± enzyme reactions were pipetted in triplicate (or more) into black 96-well plates (Corning #3603). Subsequently, 45 μL of a cocktail of glucose oxidase, horseradish peroxidase and AmplexRed (10 -acetyl-3,7-dihydroxyphenoxazine; from the components of Molecular Probes kit A22189) were added to the samples and maintained at room temperature in the dark until reading in the Synergy H1 instrument (Biotek) at 80% gain, fluorescence endpoint, excitation 530 nm, emission 590 nm. The fluorescence values from the negative controls were subtracted from experiment values to estimate the amount of glucose released from glycogen by amyloglucosidase.

Optimal establishment of primary human hepatocytes in tissue culture required a substratum of type 1 collagen (24-well plates; ThermoFisher Scientific #A11428-02) and initial plating at 200,000 cells/well in complete Williams’ E medium containing supraphysiological concentrations of insulin. This medium was essential for high-efficiency plating but precluded the study of insulin effect. We developed an insulin-free DMEM (IF-DMEM) supplemented with nicotinamide, zinc, copper, glutamine, transferrin, selenous acid and dexamethasone and no serum (37), which replaced the complete Williams’ E from Day 2 onward (after washout of insulin from the adherent monolayers). After 24 h in IF-DMEM, we re-stimulated monolayers with 5000 pM insulin (fast-acting lispro insulin; Humalog from Lilly) for 24 hr and cellular glycogen content was assessed. Glucagon (GCG) treatment was for 15 or 30 min. Glucagon-mediated glycogenolysis was complete by 15 min with no change thereafter. Each glycogen assay included a glucose standard curve (0-5 nanomoles) as an absolute reference against which we gauged the fluorescence from Resorufin in insulin or glucagon-treated primary hepatocytes and used to estimate glycogen content in nM. We also assessed the effect of siRNA knockdown of *LOC157273* on glycogen storage in hepatocytes from donor Hu8200 using the same protocol. A generalized linear model was used to estimate the mean effect of siRNA11 and siRNA15 on hepatocyte glycogen content in nM.

### Allelic imbalance of LOC157273 transcription in primary human hepatocytes

*LOC157273* allelic imbalance was measured using RNA from primary human hepatocytes from a heterozygous (A/G) donor. We estimated allele-specific *LOC157273* transcription using gene-specific strand-specific (GSSS) reverse-transcription followed by PCR (**Supplementary Text**) and analysis on EtBr-stained agarose gels. cDNA priming was performed with a gene-specific primer (**Supplementary Table S3**) targeting a region of exon 3 common to major and minor alleles. The discrimination between major (rs4841132-G) and minor (rs4841132-A) alleles relies on the subsequent PCR step utilizing the reverse primers (32A, 32G, 33A or 33G) where the most 3’-base in the PCR primer is T (for 32A or 33A) or C (for 32G or 33G)—the precise position of the SNP.

## Results

### Bioinformatic evidence that LOC157273 is a candidate causal transcript

Bioinformatic analysis showed the chr8p23.1 *PPP1R3B*-*LOC157273* locus to have the greatest amount of GWAS evidence for disease associations after intersection with all long non-coding RNA genes (**Supplementary Text, Supplementary Table S4**). The variant rs4841132 lies in a linkage disequilibrium (LD) region that spans *LOC157273* (**Supplementary Figure S2**) at a location corresponding to a promoter-like histone state in liver-derived cells (**Supplementary Table S5; Supplementary Text**). The strongest DNase I hypersensitive site lies at the conserved promoter of *LOC157273* (the gene appears to be conserved only between humans and non-human primates) and contains *CEBP* and *FOXA1* binding sites (**Supplementary Figure S3**). *LOC157273* is expressed almost exclusively in human hepatocytes (**Supplementary Figure S3**) (38). Notably, human *LOC157273* (hg19 and hg38) does not have any positional equivalents or putative orthologs in mouse (mm9 and mm10) detectable using our approaches (**Supplementary Figure S4A-B**) (39). Evident lack of conservation beyond primates is typical for human lncRNA genes (61, 62), and primate-specificity of functional lncRNAs such as *LOC157273* hints at limitations of mouse models.

### Small-interfering (si)RNA knockdown of LOC157273 reduces LOC157273 and increases PPP1R3B RNA levels and glycogen deposition in human hepatocytes

Stellaris RNA FISH in rs4841132 G/G or A/G human hepatocytes showed that *LOC157273* is a cytoplasmic lncRNA, confined to small punctate (0.5 to 1.2 micron) bodies surrounding the nuclei (**Figure 1**). As cytoplasmic lncRNAs are amenable to siRNA-mediated knockdown, in G/G hepatocytes we performed transient transfection of four different siRNAs targeting exons 1, 2, and 3 of the *LOC157273* transcript (**Figure 2A)**. The siRNAs siRNA-09 and siRNA-13 did not reproducibly reduce *LOC157273* transcript levels, but siRNA-11 and siRNA-15 reproducibly decreased *LOC157273* lncRNA by 72% (95% confidence interval (CI): 70-74%) and 75% (95% CI: 73-76%), respectively (**Figure 2B**). Knockdown with siRNA-11 and siRNA-15 increased the level of *PPP1R3B* mRNA by 57% (95% CI: 54-61%) and 79% (95% CI: 70-89%), respectively (**Figure 2C**).

**Figure 1:**
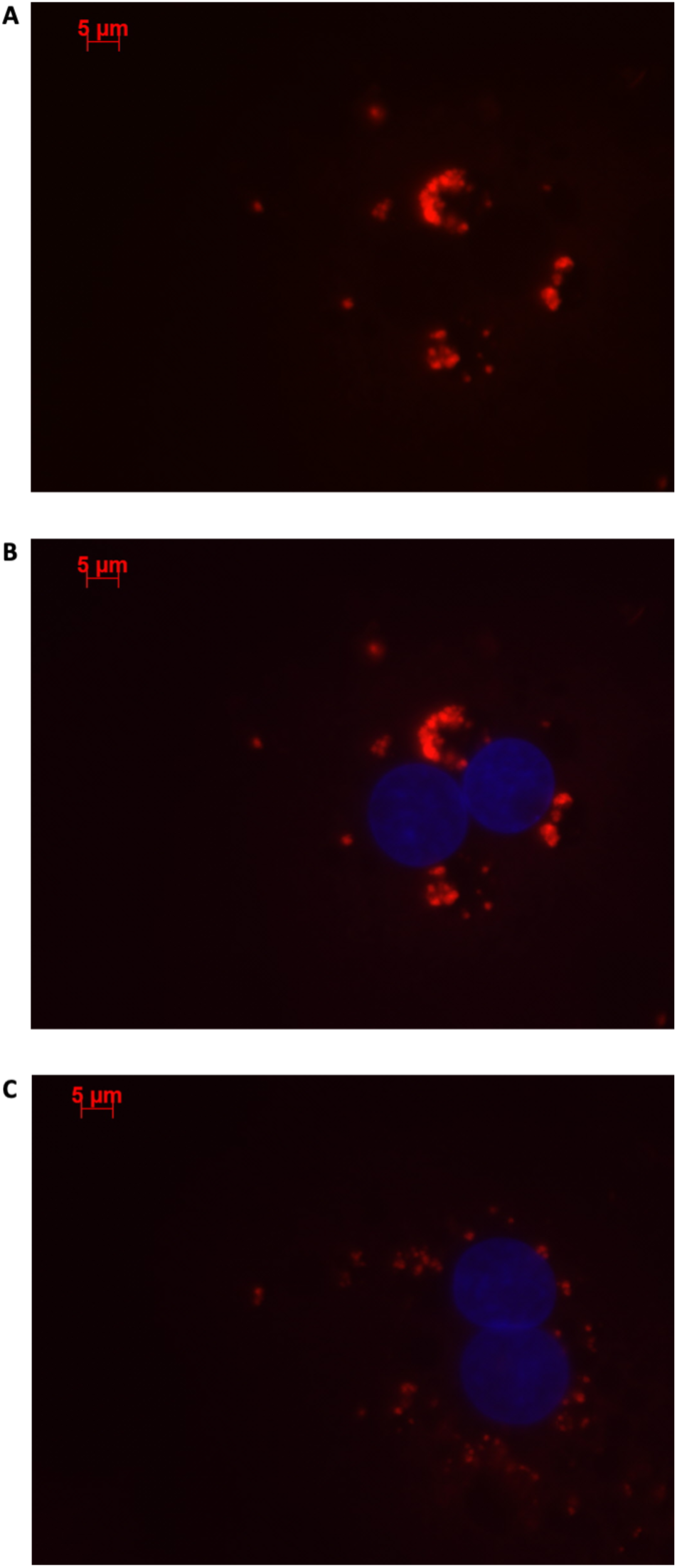
Human hepatocytes stained using Stellaris RNA FISH show that lncRNA *LOC157273* localizes to distinct punctate foci in the cytoplasm. The red staining (63x oil, 500 ms, 568 nm) in human hepatocytes with rs4841132 G/G genotype (**Panels A and B**) or with rs4841132 A/G genotype **(Panel C)** shows cytoplasmic punctate foci containing *LOC157273*, a pattern consistent with the possible localization of this lncRNA in cytoplasmic riboprotein (protein-RNA) complexes. Blue staining indicates hepatocyte nuclei.

**Figure 2:**
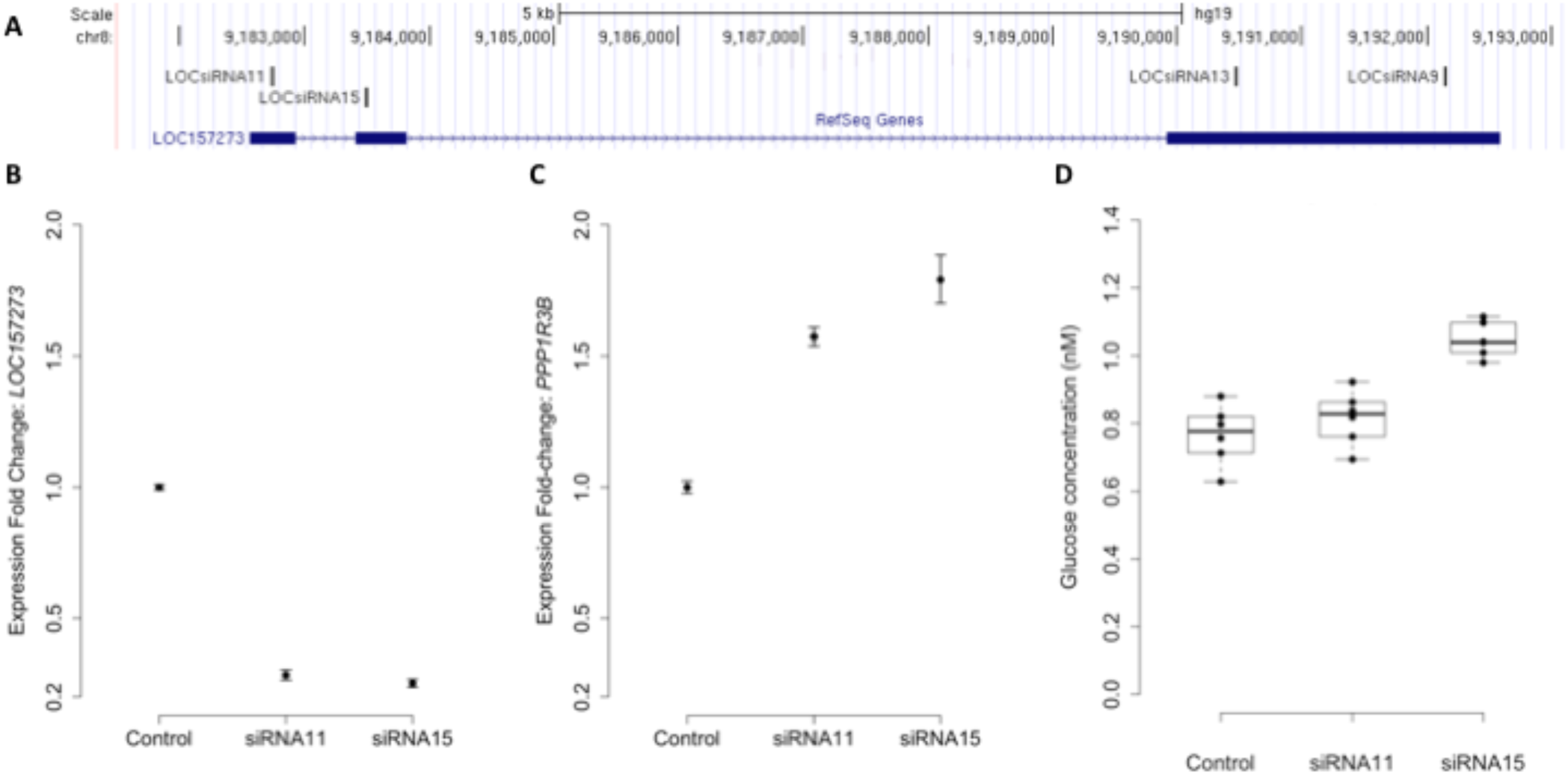
siRNA knockdown of *LOC157273* in human hepatocytes with rs4841132 G/G genotype reduces *LOC157273* lncRNA levels and increases both *PPP1R3B* mRNA levels and glycogen deposition. **Panel A** shows a UCSC Genome Browser view of the genomic position on chromosome 8 (hg19) of *LOC157273* (sense) and four siRNA constructs (antisense). siRNA-11 and siRNA-15, the most efficient constructs, targeted exons 1 and 2 of the lncRNA. **Panel B** shows expression fold-change of *LOC157273* mRNA and **Panel C**, *PPP1R3B* mRNA, after knockdown with siRNA11, siRNA15 and control. Error bars represent standard errors of normalized Taqman qRT-PCR expression normalized and averaged over 3 biological replicates. **Panel D** shows human hepatocyte glycogen content (6 replicates per condition) after knockdown with siRNA11, siRNA15 and control in one biological replicate.

We investigated the effects of *LOC157273* knockdown on cell physiology by measuring glycogen production, which is directly affected by *PPP1R3B* levels. Averaged across three biological replicates (accounting for biological variability), *LOC157273* knockdown with siRNA-15 increased insulin-stimulated glycogen deposition by 13% (P=0.002), (**Figure 2D**; **Supplementary Figure S5**). Notably, siRNA-15 is the siRNA located closest to the variant rs4841132 in exon 2 of *LOC157273*. We also tested plasmid-based overexpression of *LOC157273* in human hepatocytes and hepatoma cells but did not find alteration in levels of *PPP1R3B* expression (**Supplementary Text, Supplementary Figure S6**).

### As expected, LOC157273 knockdown has diverse transcriptome-wide effects

We performed a gene expression differential expression analysis of human hepatocyte whole transcriptomes comparing siRNA-11 and siRNA-15 knockdown to control conditions. Of 15,441 unique genes tested, 953 genes showed nominal evidence for differential expression at P < 0.01 **(Figure 3A, Supplementary Table S6**). RNA-seq results were consistent with Taqman RT-PCR results for both *LOC157273* (Fold Change = 0.75, P= 0.03) and *PPP1R3B* (Fold Change = 1.34, P = 0.001). To elucidate biological pathways which might be affected by these expression changes, we performed a Reactome pathway analysis with three sets of genes: the 953 genes with P<0.01 (**Supplementary Table S7A**), the 206 genes with Fold Change ≤ 0.74 and P_adj_ < 0.001, and the 222 genes with Fold-change ≥ 1.34 and P_adj_ < 0.001 (**Supplementary Figure S7; Supplementary Table S7B**). We observed two nominally significant Reactome pathways with the genes with Fold Change ≤ 0.74 and P_adj_ < 0.001: Glucuronidation and Biological Oxidations (P=0.04 for both pathways), and 17 enriched Reactome pathways with the genes with Fold Change ≥ 1.34 and P_adj_ < 0.001 (**Figure 3B)**. An unbiased gene-set enrichment analysis, using all gene results from the differential expression analysis, showed 164 enriched pathways with P<0.05, including fatty acid metabolism (P=0.0004), glucuronidation (P=0.0007), gluconeogenesis (P=0.02), and glucose metabolism (P=0.03) (**Supplementary Table S8**). Of the significantly enriched pathways, 23% fall under ‘Cell Cycle’ in the Reactome hierarchy, 16% under ‘Metabolism’ and 12% under ‘Signal Transduction.’

**Figure 3:**
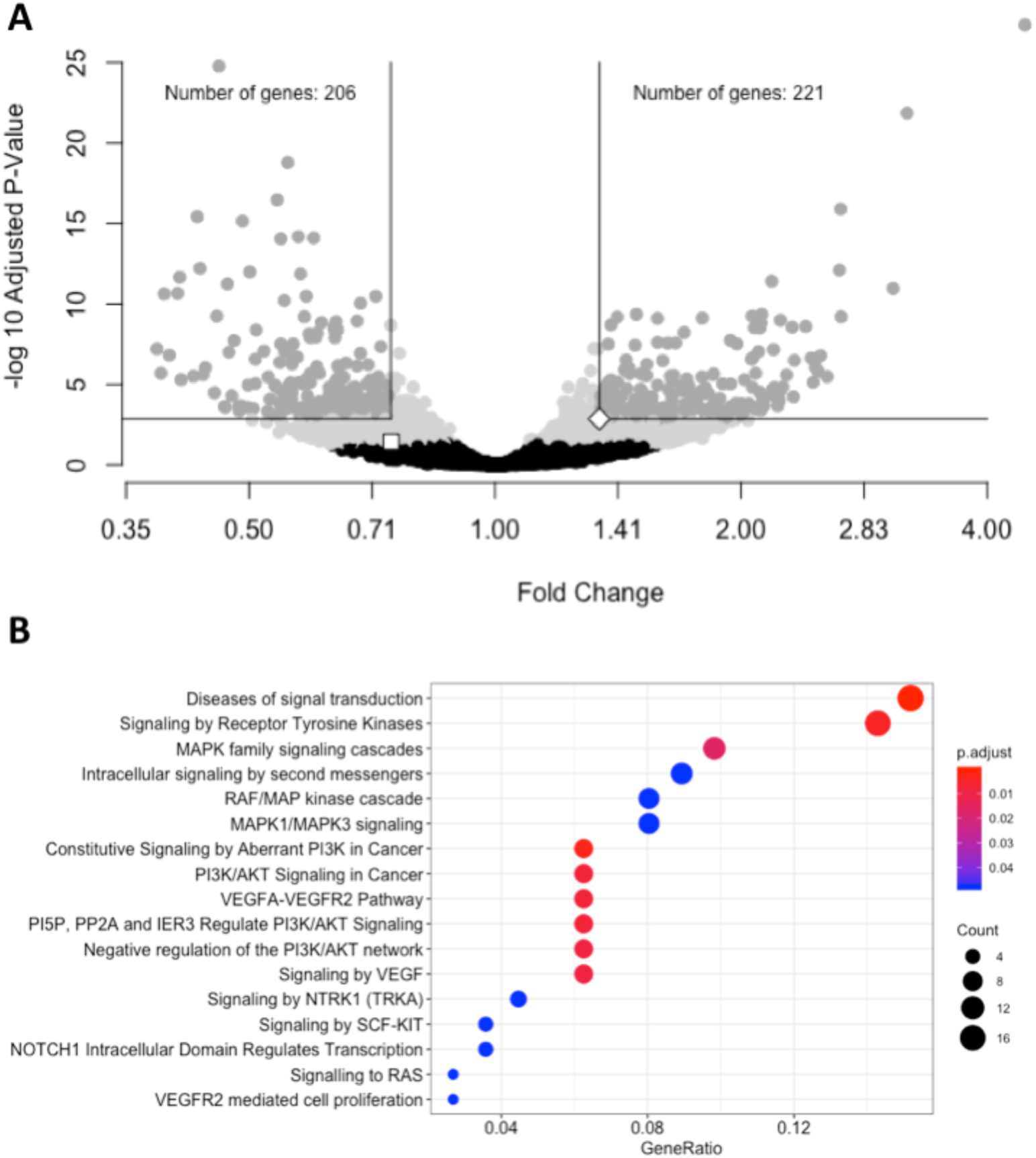
siRNA knockdown of *LOC157273* using transcriptome-wide differential expression analysis in rs4841132 G/G genotype human hepatocytes results in significant expression changes in hundreds of transcripts, including *LOC157273, PPP1R3B* and numerous putative *trans* targets. **Panel A.** Volcano plot showing the fold-change effect of siRNA-11 and siRNA-15 knockdowns combined (X axis), compared with controls, with gray points indicating significant change in transcript expression (family-wise P_adj_ < 0.05) (Y axis). The white square point indicates the *LOC157273* transcript and shows that its knockdown reduced its expression (fold change = 0.74, P = 0.004, P_adj_ = 0.04), as expected for a successful knockdown experiment. The white diamond point indicates the *PPP1R3B* transcript and shows that *LOC157273* knockdown increased *PPP1R3B* expression 34% (fold change = 1.34, P = 4.5 x 10^−5^, P_adj_=0.001). **Panel B.** Reactome enrichment analysis of genes with increased expression after siRNA-11 or siRNA-15 knockdown compared to controls (221 dark gray points in Panel A; Fold Change > 1.34 and P_adj_ < 0.001). Each row represents a significant Reactome pathway (family-wise P < 0.05) with GeneRatio (X axis) showing the degree to which the differentially expressed genes were enriched in the pathway. The count of differentially expressed genes within each pathway is depicted with the size of the circles, and the significance of the enrichment is depicted with the color of the circles.

### Glycogen content and allelic imbalance of LOC157273 transcription in human hepatocytes

Patterns of *LOC157273* expression and glycogen deposition in the rs4841132 A/G carrier were similar to those seen by siRNA knockdown of *LOC157273*: ΔCT comparing *LOC157273* to *GAPDH* for G/G carriers averaged 8.94 (95% CI: 8.86-9.01), while the A/G carrier showed less *LOC157273* lncRNA (ΔCT: 11.0 (95% CI: 9.4-12.6), an estimated 76% decrease in expression in A versus G allele carriers (**Table 1**). Before insulin stimulation, the median glycogen store in hepatocytes from the A/G heterozygote was 0.40 nM in a first replicate and 0.44 nM in a second replicate, compared to 0.12 nM in hepatocytes from a G/G homozygote (**Figure 4**). Median basal glycogen concentration in two other G/G donors were 0.14 nM and 0.06 nM (**Supplementary Figure S8**). In one assay, A/G hepatocytes had median glucose concentration of 0.53 nM after insulin re-stimulation, representing a 31% increase (P=0.004) in glycogen content over basal levels. In a replicate, glucose concentration was 0.59 nM after insulin re-stimulation, representing a consistent but non-significant increase of 33% (P=0.06). The hepatocytes from the G/G donor had no observable increase (P=0.7) in glycogen content over basal levels (**Figure 4)**. To test whether the G or the A allele was responsible for the observed effects on *LOC157273* action, we used a single reverse primer in exon 3 (**Supplementary Table S3**) to prime cDNA synthesis enriched for *LOC157273* cDNA. In one assay, we observed 78% reduction of the rs4841132-A (minor) allele transcript compared to the G allele transcript. In a replicate, we observed 88% reduction (**Figure 5**). These results suggest that the diminished content of *LOC157273* lncRNA in this A/G heterozygous hepatocyte donor results from a specific loss of lncRNA output in rs4841132-A (minor) allele carriers.

**Table 1:** Difference in CT (ΔCT) for target gene and *GAPDH* reference gene from qRT-PCR on RNA purified from primary human hepatocytes derived from four homozygous (G/G) donors and one heterozygous (A/G) donor

| Donor | Replicates | Genotype | ΔCT for LOC157273<br>Mean (Min–Max; SD) | ΔCT for General-PPP1R3B<br>Mean (Min–Max; SD) | ΔCT for Hepatocyte-<br>specificPPP1R3B<br>Mean (Min–Max; SD) |
| --- | --- | --- | --- | --- | --- |
| Hu8200A* | 4 | G/G | 8.7 (5.1 – 11.5; 2.7) | 8.0 (7.1 – 8.7; 0.8) | 9.0 (8.0 – 10.0; 1.0) |
| TRL4056B | 3 | G/G | 9.5 (8.8 – 9.8; 0.6) | 6.8 (5.2 – 7.6; 1.4) | 4.5 (3.8 – 5.0; 0.6) |
| TRL4105A | 4 | G/G | 8.5 (8.0 – 9.2; 0.6) | 7.6 (5.9 – 9.3; 1.8) | 7.6 (5.6 – 9.0; 1.6) |
| TRL4108 | 3 | G/G | 8.7 (7.8 – 9.3; 0.8) | 7.3 (6.0 – 7.9; 1.1) | 7.2 (6.5 – 7.8; 0.7) |
| **Meta-Analysis of 4 G/G donors** |  |  | **8.94 (SD: 0.131)** | **7.52 (SD: 0.311)** | **6.39 (SD: 0.153)** |
| TRL4079 | 4 | A/G | 11.0 (10.2 – 12.4; 0.9) | 7.7 (5.1 – 10.2; 2.9) | 6.9 (5.5 – 8.4; 1.5) |

**Figure 4:**
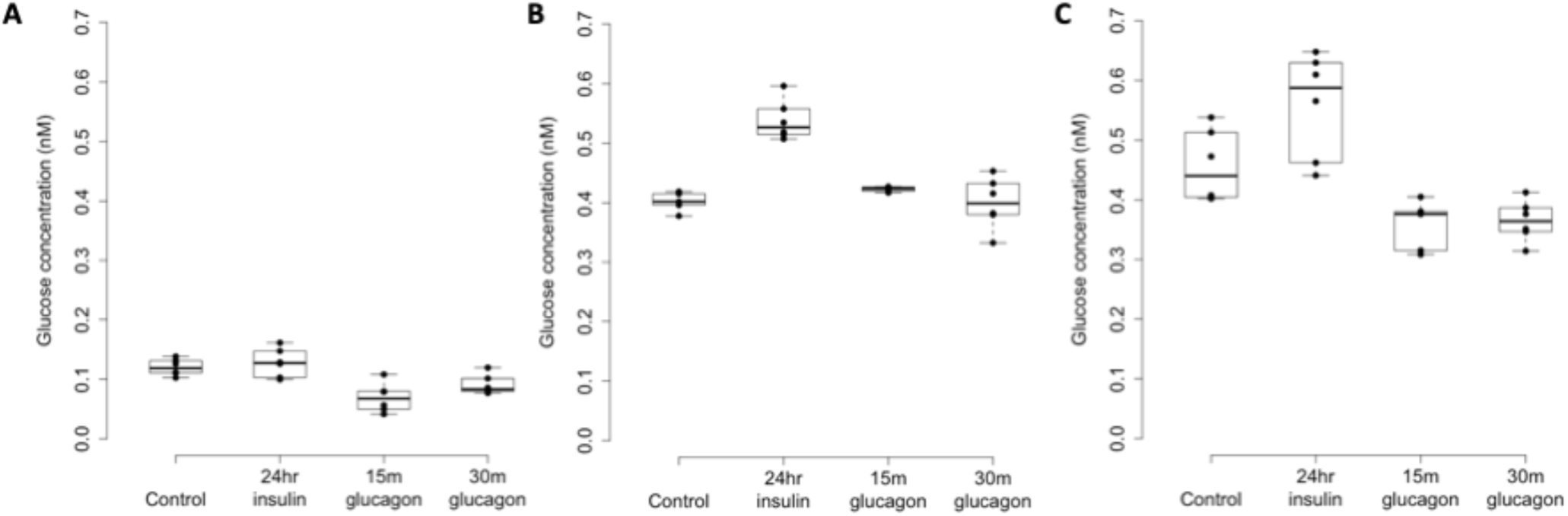
Glycogen deposition response after insulin re-stimulation of primary human hepatocytes. **Panel A** shows a donor with rs4841132 G/G genotype and **Panels B and C** show replicates of a donor with rs4841132 A/G genotype. Brief (15 or 30 min) treatment with 5 nM glucagon demonstrates a decrease in glycogen compared to the control, confirming that what is being measured is glycogen.

**Figure 5:**
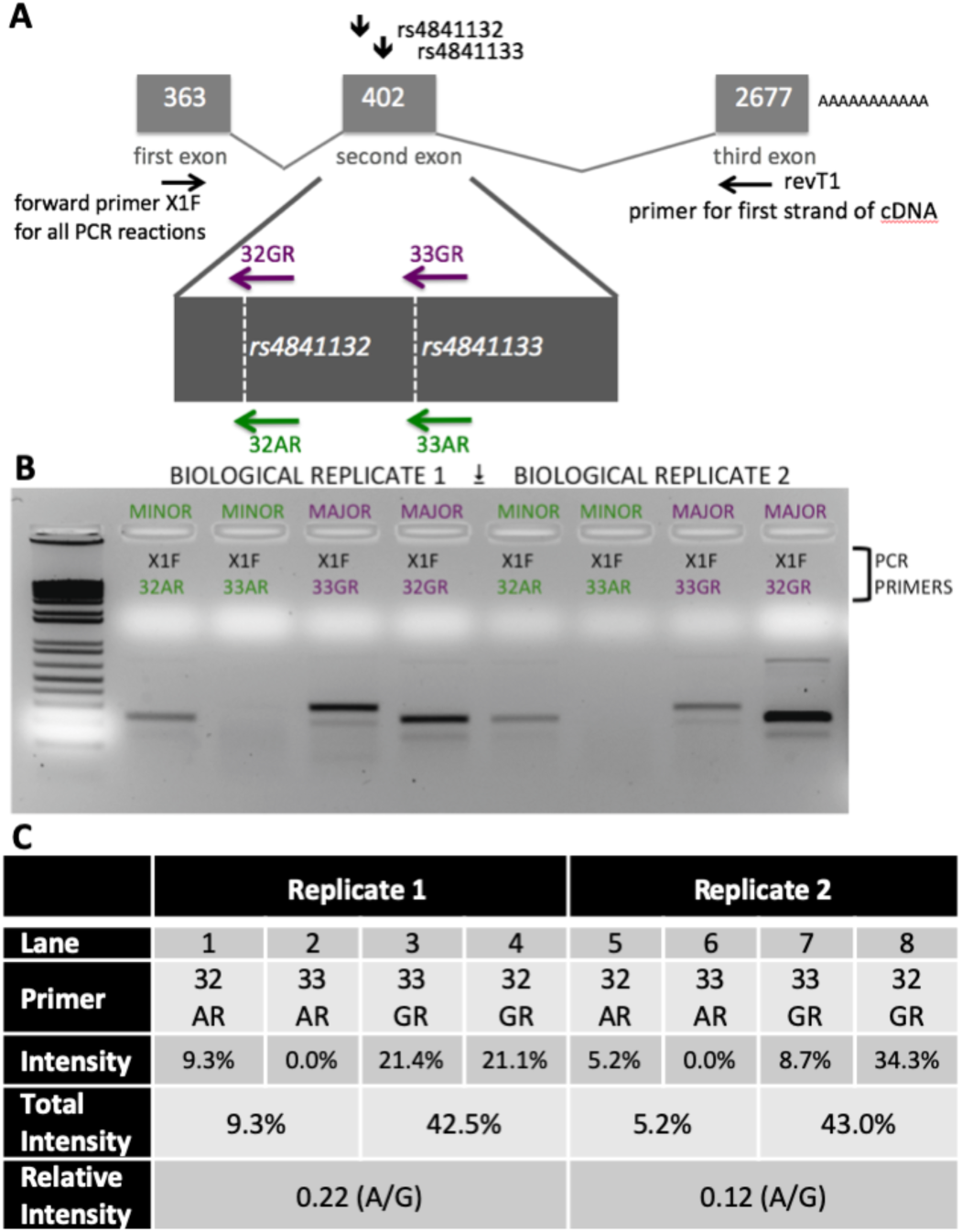
Analysis of *LOC157273* transcription in primary hepatocytes from a heterozygous rs4841132 A/G donor demonstrates decreased transcription of the minor (A) allele. **Panel A.** RNA was transcribed into cDNA using a gene-specific strand-specific primer for two replicates. **Panel B.** Aliquots of cDNA were amplified with PCR primers seated in the first exon (black, X1F) and allele-specific primers that recognize only the major allele (magenta, 32G or 33G) or minor allele (green, 32A or 33A). he X1F-32 reaction yields a PCR product of 231 bp, while the X1F-33 reaction yields a PCR product of 302 bp. **Panel C.** Image-based quantification of the X1F-32 and X1F-33 bands from the minor allele compared to the major allele using the ImageJ software. Total intensity: sum of 32GR/33GR or 33AR/32AR lanes. Relative Intensity: quantification of the rs4841132-A minor allele intensity compared to the rs4841132-G major allele intensity.

## Discussion

Over fifteen years of genome-wide association discovery provides a rich abundance of loci and variants associated with T2D risk and its underlying metabolic pathophysiology. The genetic architecture of T2D is dominated by non-coding, regulatory, mostly common variation; with only ∼7% of variants convincingly shown to directly encode protein-altering mutations in Europeans (6, 8). The function of these hundreds of non-coding variants is being illuminated by integration of genomic association data with genomic functional information; for T2D, most data thus far come from islet chromatin regulatory mapping (40-42). Islet genomic regulatory features linked to GWAS signals include, for instance, enhancer site disruption (43), stretch enhancer potentiation (44), and lncRNA action in human (45) and mouse beta cell lines (25, 46).

Liver, muscle and fat tissue also contribute to T2D pathophysiology, where progress is also being made to link human tissue-specific regulatory maps to GWAS signals (47, 48). In this report, we present evidence in human liver cells that the lncRNA *LOC157273* is the causal transcript at the GWAS-discovered chr8p23.1 “*PPP1R3B*” locus. We show that *LOC157273* is expressed exclusively in human hepatocytes, is a close (< 200 kb) genomic neighbor of an attractive T2D physiologic candidate gene, *PPP1R3B*, and is a negative regulator of *PPP1R3B* expression.

We demonstrated that *LOC157273* is a predominantly cytoplasmic lncRNA. Any roles it may play in the cytoplasm remains uncertain. It is unknown whether a cytoplasmic mechanism exists whereby which *LOC157273* may regulate its neighbor gene *PPP1R3B* (presumably independent of genomic proximity). We also cannot discount an alternative role for *LOC157273* in genomic regulation, nor a potential role for *LOC157273* in dual control in both cytoplasmic and nuclear regulation of *PPP1R3B*. For instance, *LOC157273* might be subject to bidirectional nucleocytoplasmic shuttling with cytoplasmic excess, but still has a nuclear function in epigenetically downregulating its neighbor gene. If true, this would explain both the cytoplasmic foci (putative RNA-protein complexes allowing *LOC157273* to regulate numerous genes in-trans) and why its cytoplasmic knockdown rescues *PPP1R3B* expression (removal of excess *LOC157273* which cannot go back to the nucleus to regulate its neighbor gene in-cis).

SiRNA knockdown of *LOC157273* nearly doubled *PPP1R3B* mRNA levels (versus control) and altered the expression of other human hepatocyte transcripts. This was accompanied by an increase of over >50% in insulin-mediated hepatocyte glycogen deposition, which is an expected functional consequence of increased *PPP1R3B* expression. As PPP1R3B is the principal known regulator of glucose entry into the hepatic glycogen deposition pathway, its negative regulation by *LOC157273* (containing rs4841132) strongly supports the lncRNA and not the protein *per se* as the causal transcript at the locus. Indeed, evidence for common genetic variation in *PPP1R3B* itself has been inconsistent for T2D risk in humans (21, 49), although rare variants in *PPP1R3B* are associated with human diabetes phenotypes (50, 51).

Negative regulation of *PPP1R3B* by *LOC157273* provides partial evidence for the lncRNA to be the functional transcript at the chr8p23.1 locus. As the rs4841132-A allele occurs in about 1 in 10 people, we were able to find a single rs4841132 A/G heterozygote among all the commercial hepatocyte donors available to us. This A/G carrier had reduced *LOC157273* abundance, increased *PPP1R3B* expression and increased glycogen deposition vs. averages from three G/G carriers, mimicking the effect observed with knockdown in G/G hepatocytes. Although heterozygote data are from a single donor, they are concordant with other, independent data. In 125 liver biopsy samples from obese patients, the rs4841132-A allele (vs G) was associated with higher *PPP1R3B* mRNA levels, reduced LOC157273 expression and protection against histologic hepatic steatosis (29). In another cohort of 1,539 individuals with non-viral liver disease, the rs4841132-A allele (vs G) was associated with increased hepatic x-ray attenuation reflecting increased glycogen deposition, consistent with a mild form of hepatic glycogenosis (30). Our data suggest that the rs4841132-A allele confers decreased transcriptional efficiency of *LOC157273*; reduced lncRNA abundance induces *PPP1R3B* upregulation as well as other transcripts apparently related to increased glycogen deposition. The large number of genes with differential expression after *LOC157273* knockdown (221 upregulated and 206 downregulated) support a scenario of *LOC157273* as a master regulator of transcription in *trans*. Furthermore, several enriched pathways (fatty acid metabolism, PI3K/AKT signaling, among others) are concordant with enriched pathways from the transcriptome analysis of the 125 liver biopsy samples comparing carriers of the rs4841132-A allele vs non-carriers (29). Taken together, the functional data we present support the contention that the lncRNA *LOC157273* is the causal transcript at the chr8p23.1 GWAS locus, controlling T2D risk and metabolic physiology by hepatic regulation of *PPP1R3B* (and most likely other) transcription, thereby influencing variation in glycemia, other metabolic phenotypes, and T2D risk observed in genetic association studies.

LncRNAs as a class have diverse molecular functions, and a few lncRNAs have emerged to be associated with cardiometabolic disease (52). *MIAT* (myocardial infarction associated transcript) was the first lncRNA identified by GWAS as a disease candidate gene (22). The chromosome 9p21 lncRNA *ANRIL* was subsequently shown to be associated with several forms of atherosclerosis; we now know that this molecule confers protection from atherosclerosis by controlling ribosomal RNA maturation and modulating atherogenic molecular networks (53). The regional chr9p21 GWAS signal has also been associated with T2D, but this association is not mediated by *ANRIL*, as the association signal for T2D is separated from that for atherosclerosis by a recombination hot spot that renders the two disease association signals in linkage equilibrium (54). Human pancreatic islets transcribe thousands of lncRNAs, many of which are highly islet- or beta cell-specific (25), including two beta cell lncRNAs shown to cis-regulate nearby genes involved in T2D physiology. Loss-of-function screening in a human islet beta cell line identified the lncRNA *PLUTO* (PDX1 locus upstream transcript) as a positive regulator of *PDX1*, a critical transcriptional regulator of human pancreas development and beta cell function (45). *PLUTO* appears to cis-regulate *PDX1* by altering chromatin structure to facilitate contact between the *PDX1* promoter and its enhancer cluster. Another islet-specific transcript, *blinc1*, has been shown to regulate the expression of groups of functionally related genes, including *NKX2-2*, an essential transcription factor important for beta cell developmental programs (46).

Integrating these observations with prior evidence, we envision the following physiological model for the action of *LOC157273* on T2D hepatic physiology (**Supplementary Figure S9)**. Our siRNA data suggest that *LOC157273* is a functional suppressor of *PPP1R3B* transcription, with lower lncRNA levels associated with higher hepatocyte *PPP1R3B* expression. PPP1R3B is the glycogen-targeting subunit of phosphatase PP1, regulating PP1 activity by suppressing the rate at which PP1 inactivates glycogen phosphorylase (decreasing glycogen breakdown) and enhancing the rate at which it activates glycogen synthase and increases glycogen synthesis (55). Increased liver *PPP1R3B* expression therefore shifts basal and insulin-stimulated hepatic glycogen flux towards storage (16), which is consistent with our *in vitro* data as well as human liver imaging studies showing that the rs4841132-A allele carriers have increased hepatic attenuation on CT imaging, suggestive of increased glycogen storage (15, 56). Assuming that rs4841132-A allele carriers have increased glycogen stores, we hypothesize that these will lead to reduced glucose-uptake by the liver in the fasting state, consistent with observed GWAS associations with elevated fasting serum glucose and insulin levels and T2D risk (4). Additionally, abundant liver glycogen in the fasting state may be easily mobilized to glucose-6-phosphate and therefore increase glycolytic flux, consistent with observed associations with increased lactate levels in humans (57). Liver free fatty acid formation is also influenced, consistent with observed decreases in cholesterol levels (58). In contrast to the fasting state, in the postprandial state, the rs4841132-A allele appears to cause increased insulin-mediated glucose-uptake the by the liver (16). With liver glycogen synthesis increased, hepatic glucose uptake may increase, consistent with observed associations for SNPs near *PPP1R3B* with decreased 2-hour post OGTT glucose levels seen in our earlier GWAS (4).

Strengths of our study include a clear test of the hypothesis that *LOC157273*, in which the well-documented metabolic trait variant rs4841132 resides, regulates the nearby gene *PPP1R3B* and influences hepatocyte glycogen deposition, supporting the contention that the lncRNA is the causal transcript at the chr8p23.1 GWAS locus. That we demonstrated this in humans is important, as *LOC157273* does not have an orthologue in rodents and can’t be meaningfully studied in them. Limitations include that we studied allelic imbalance in only one rs4841132 A/G heterozygote hepatocyte donor, but effects in this individual were similar to lipid and glycogen hepatic storage effects seen in much larger samples with A/G carriers. Unbiased RNA-seq following siRNA knockdown of *LOC157273* identified many additional transcripts that could contribute to the observed increase in hepatic glycogen deposition, but due to the small number of biological replicates available, full exploration of these signals will require future research. Finally, we do not yet know the exact molecular mechanism whereby *LOC157273* regulates *PPP1R3B* or any of the other transcripts seen by RNA-seq. LncRNAs have multiple mechanisms that influence gene regulation; here, we might postulate epigenetic and/or cytoplasmic mechanisms to explain both its cytoplasmic location and apparently broad transcriptional effects (59).

Causal transcripts, relevant tissues and molecular mechanisms underlying the hundreds of T2D-associated genomic loci are now coming to light. A goal of modern chronic disease genomics is to identify new mechanisms for therapeutic targeting. RNA therapeutics is one novel frontier for genomic medicine. Small-interfering (si) RNA therapeutics, which as in our approach loads its target RNA into the endogenous cytoplasmic RNA-induced silencing complex (RISC), are now in late-stage clinical trials for lipid-lowering through PCSK9 inhibition (60). The liver is especially amenable to RNA therapeutics; glycemic control via siRNA regulation of glycogen entry into the liver arises as a tantalizing possibility. However, the mild glycogenosis that appears to accompany genetic variation at rs4841132 raises the possibility that lowering blood glucose by increasing hepatic storage of glycogen may have its own risks (30). While our data support a genetic regulatory relationship between *LOC157273* and *PPP1R3B, LOC157273* appears to regulate many other transcripts whose action in regulating hepatic glycogen and cholesterol flux remain to be explained. Nonetheless, we have identified a lncRNA to be the causal transcript at a hepatic T2D-cardiometabolic disease GWAS locus, opening the window to the possibility of new, RNA-based therapeutic pathways for therapy and prevention of T2D.

## Acknowledgements

Funding: AKM was supported by K01 DK107836. RS supported by U01 DK078616. JBM supported by R01 DK078616, U01 DK078616, K24 DK080140, and an American Diabetes Association Mentored Scientist Career Development Award. LL supported by the NIH Director’s New Innovator Award 1DP2-CA196375, and by the Wayne State University 2018-2019 Charles H. Gershenson Distinguished Faculty Fellowship. MG supported by the Eris M. Field Chair in Diabetes Research. JR: supported in part by R01 DK078616, the National Center for Advancing Translational Sciences, CTSI grant ULTR001881, and the National Institute of Diabetes and Digestive and Kidney Disease Diabetes Research Center (DRC) grant DK063491 to the Southern California Diabetes Endocrinology Research Center. MG and JR supported by Diabetes Research Center grant P30 DK063491 and National Center for Advancing Translational Sciences (NCATS) Grant UL1TR001881

JBM is the guarantor of this work and, as such, had full access to all the data in the study and takes responsibility for the integrity of the data and the accuracy of the data analysis.

## Author Contributions

Led the study: JM, LL; Writing initial draft and submitted final draft of paper: AM, AG, EK, RS, JM, LL; Interpreted data, provided intellectual input, reviewed drafts of the paper and approved the final version: All; Conducted laboratory experiments: AG, PT, JC, JD; Conducted statistical analysis and bioinformatics: AM, EK, JB; Provided funding and material support: JM, LL

**The authors declare no conflict of interest.**

