## Supplementary Text and Figures for "A long noncoding RNA, *LOC157273*, is the effector transcript at the chromosome 8p23.1-*PPP1R3B* metabolic traits and type 2 diabetes risk locus"

**Institutions**

1. Clinical and Translational Epidemiology Unit, Mongan Institute, Massachusetts General Hospital, Boston, MA, USA
2. Department of Medicine, Harvard Medical School, Boston, MA, USA
3. Programs in Metabolism and Medical & Population Genetics, Broad Institute of MIT and Harvard, Cambridge, MA, USA
4. Center for Molecular Medicine & Genetics, Wayne State University, Detroit, MI, USA
5. Karmanos Cancer Institute at Wayne State University, Detroit, MI, USA
6. Center for Human Genetics Research, Massachusetts General Hospital, Boston, MA, USA
7. Division of General Internal Medicine, Massachusetts General Hospital, Boston, MA, USA
8. Diabetes Unit, Massachusetts General Hospital, Boston, MA, USA
9. Department of Statistics, University of California, Berkeley, CA, USA
10. Centre for Computational Biology, University of Birmingham, B15 2TT Birmingham, United Kingdom.
11. Division of Endocrinology, Diabetes, and Metabolism, Department of Medicine, Cedars-Sinai Medical Center, Los Angeles, CA, USA
12. The Institute for Translational Genomics and Population Sciences, Departments of Pediatrics and Medicine, Los Angeles Biomedical Research Institute at Harbor-UCLA Medical Center, Torrance, CA USA
13. Department of Human Genetics, McGill University, Montréal, Québec, Canada.
14. Department of Medicine, McGill University, Montréal, Québec, Canada
15. McGill University and Genome Québec Innovation Centre, Montréal, Québec, Canada.
16. Department of Neurology, School of Medicine, Wayne State University, Detroit, MI, USA

* Contributed equally

### Supplement Table of Contents

***Supplementary Text*** **3**
Supplementary material providing additional detail on methods: 3
Cellular localization of LOC157273 with Stellaris RNA fluorescent in situ hybridization 3
TaqMan quantitative reverse-transcriptase PCR (qRT-PCR) 3
Allelic imbalance of LOC157273 transcription 3
Human hepatocyte sources and culture 4
Screen of human hepatocyte donors to identify rs4841132 genotype 4
Bioinformatic evidence that *LOC157273* is a candidate causal transcript 5
*LOC157273* conservation 5
Alternative quantification of allele-specific *LOC157273* transcription using cDNA cloning 6
Targeted resequencing of 5.6 kb of genomic DNA covering the gene body and upstream region of *LOC157273* in a single heterozygote hepatocyte donor 6
Plasmid-based overexpression of LOC157273 cDNA in primary human hepatocytes and hepatoma cells 7
***Supplementary Figures*** **8**
Supplementary Figure 1: Screen of the LOC157273 region from 16 human hepatocyte donors to identify rs4841132 genotype 8
Supplementary Figure 1b: DNA resequencing of major and minor alleles of the LOC157273 shows heterozygosity at SNP rs2169387 (-1170 relative to the transcriptional start site), a SNP associated with serum total cholesterol. 8
Supplementary Figure 1c: Sanger sequence reads for a LOC157273 heterozygote hepatocyte donor show half-height A/G peaks at both SNP rs4841132 and rs4841133. 9
Supplementary Figure 2: Genetic associations with fasting glucose and fasting insulin at the chromosome 8 PPP1R3B-LOC157273 locus in the MAGIC, AAGILE and CHARGE consortia. 10
Supplementary Figure 3: Expression profile of *LOC157273* 12
Supplementary Figure 4: Comparative genomics of *LOC157273* in modern and ancient humans, nonhuman primates, and mouse. 14
Supplementary Figure 5: Glycogen content in human hepatocytes (Hu8200 donor homozygote G/G for rs4841132) in siRNA knockdown vs siRNA control conditions. 17
Supplementary Figure 6: Fold-change in *LOC157273* and *PPP1R3B* expression after transfection with plasmid to overexpress *LOC157273* 18
Supplementary Figure 7. Cluster analysis showing clustered genes with similar normalized transcript counts in siRNA11, siRNA15 and control conditions from the transcriptome-wide differential expression analysis. 19
Supplementary Figure 8: Insulin re-stimulation of primary human hepatocytes stimulates glycogen deposition in hepatocytes from additional heterozygous (G/G) donors. 20
Supplementary Figure 9: Physiological Model of *LOC157273* action on *PPP1R3B*, hepatic glycogen flux and peripheral glucose and insulin levels 21
***Supplementary Tables*** **22**
Supplementary Table 1: Primary Human Hepatocyte Donor Characteristics 22
Supplementary Table 2. TaqMan primers chosen for two PPP1R3B mRNA known isoforms and a single set for LOC157273 23
Supplementary Table 3: Primers used in allelic-imbalance of LOC157273 transcription in a heterozygous hepatocyte donor 24
Supplementary Table 4: Genomic locations after intersection of GENCODE v18 exons with GWAS SNPs 25
The following Supplementary tables are in Supplement_LOC-PPP_Paper_05May2019.xlsx 27
Supplementary Table 5. Haploreg (v4) query of rs4841132. 27
Supplementary Table 6: Genes with Padj<0.01 from gene expression differential expression analysis of the whole transcriptome comparing siRNA-11 and siRNA-15 to control. 27
Supplementary Table 7: Enriched Reactome pathways from the genes showing increased expression in siRNA11 and siRNA15 vs control. 27
Supplementary Table 8: Reactome pathways showing enrichment in differential expression from Gene Set Enrichment Analysis. 27
Supplementary Table 9: Primers used to re-sequence cloned *LOC157273* genomic amplicons 28
***Supplementary References*** **29**

### Supplementary Text

#### Supplementary material providing additional detail on methods:

##### Cellular localization of LOC157273 with Stellaris RNA fluorescent in situ hybridization

We used a custom-synthesized 48-probe set (LGC Biosearch Technologies; Petaluma, CA) of non-overlapping fluorescent-tagged oligonucleotides that tiled along the 3.4 kb long non-coding RNA LOC157273. The probe-set was aligned to the GRCh37/hg19 reference genome using BLAT and the UCSC Genome Browser with the total coverage spanning 10300 bp of the reference genome, including 6 single-nucleotide variants (one in exon 1 (rs9650619, $r^2$=0.1 with rs4841132 across human populations (1, 2) two in exon 2 (G at rs4841132 and G at rs4841133, $r^2$=0.8) and three in exon 3 (G at rs7003905, $r^2$=0.1 and G at rs7004248, $r^2$=0.2)], while the majority of the probes map to the third exon (2.7 kb) and two probes (5 and 13) span introns 1 and 2 respectively. The probeset was stored at -20°C as a pool of 48 resuspended in TE buffer and kept in the dark when thawed. To prepare coverslips, primary hepatocytes were seeded on collagen type 1-coated round glass coverslips (25 mm Ø, Fisher Scientific #12-545-86) in complete Williams' E medium and cultured for 16-32 hours before fixation in 4% paraformaldehyde (1X PBS, 20 min, RT) and 2x PBS washes. Coverslips were stored in the dark at -20°C in 70% ethanol until hybridization, following the Biosearch Technologies Stellaris FISH protocol for adherent cells (https://www.biosearchtech.com/support/resources/stellaris-protocols). Probesets were hybridized with seeded coverslips for sixteen hours at 37°C in formamide-containing buffer (Biosearch Technologies #SMF-HB1-10) and then coverslips were washed to remove unbound probe. Coverslips were mounted onto glass microscope slides (25 x 75 mm) using Vectashield with DAPI Mounting Medium (Vector Laboratories #H1200) and examined under the AxioObserver inverted fluorescence microscope (Carl Zeiss Microscopy) equipped with objective lens 63×/1.40 oil. The 365 nm light-emitting diode (LED) was used to image the DAPI fluorophore; the 555 nm LED was used to image the Quasar 570 fluorophore. Images were captured using the AxioCam MRm camera (Carl Zeiss Microscopy). Red bodies in the merged images denote the Quasar 570 signal from lncRNA molecules; the blue-colorized DAPI staining shows cell nuclei.

##### TaqMan quantitative reverse-transcriptase PCR (qRT-PCR)

For all TaqMan qRT-PCR analyses, we transcribed RNA into cDNA with oligo(dT) priming with Superscript III reverse transcriptase (ThermoFisher Scientific #18080093). Aliquots of cDNA served as input for PCR reactions targeting GAPDH, LOC157273, and two isoforms of PPP1R3B. Primers to estimate PPP1R3B mRNA were designed to target the respective mRNAs of the two isoforms of PPP1R3B (Ensembl transcripts ENST00000310455.3 and ENST00000519699.1). Primers and probesets for all cDNAs spanned the exon-exon junctions (**Supplementary Table 2**). Primers and probesets were custom-ordered from Life Technologies/ThermoFisher Scientific. The amplicons were validated by cloning the final TaqMan product in the TA Cloning System (vector pCR2.1TOPO) and sequencing the products. The LOC157273 amplicon spans intron 1 and includes 36 nt of LOC157273 exon 1 and 41 nt of exon 2. Real-time qRT-PCR reactions were all done in 96-well plates (ThermoFisher Scientific #4346906), using Roche Faststart Universal Probe Master (Rox; Sigma-Aldrich Roche #4914058001) monitored on the Applied Biosystems 7500 Fast Real-Time Instrument.

##### Allelic imbalance of LOC157273 transcription

To investigate allelic imbalance in LOC157273 transcription in primary human hepatocytes from a heterozygous (A/G) donor, we estimated transcription of the major and minor LOC157273 allele RNA using gene-specific strand-specific (GSSS) reverse-transcription followed by PCR [GSSSRTPCR; (63)] and analysis on EtBr-stained agarose gels. Complementary DNA priming was done with a gene-specific primer (**Supplementary Table 3**) targeting a region of exon 3 common to major and minor alleles. The discrimination between major and minor relies on the subsequent PCR step, where we design reverse primers (32A, 32G,

33A or 33G) where the most 3'-base in the PCR primer is T (for 32A or 33A) or C (for 32G or 33G)—the precise position of the SNP. We primed 5 µg total RNA (donor TRL4079) with the gene-specific (GS) primer LOC157273_revTprimer1 located in exon 3 (5'-GGAAGACCACCAAAGAGATTATTC-3'OH) (**Supplementary Table 3**). The primer for reverse transcription lies downstream of rs4841133. The cDNA synthesis reaction was primed at 70°C for 5 min (RNA, GSSS primer, dNTPs) and then placed on ice. To the primed reaction (10 µL), we add a 10 µL cocktail of 10X RT buffer (2 µL), 25 mM magnesium chloride (4 µL), 0.1 M dithiothreitol (DTT; 2 µL), RNAse OUT (40 U/µL; 1 µL; ThermoFisher Scientific # 10777019), SuperScript III (Invitrogen; 1 µL of 200 U/µL). cDNA synthesis was done at 50°C for 50 min (SuperScript III), terminated by heating to 85°C (5 min), followed by chilling on ice. To degrade the RNA, 1 µL of RNAse H was added to the cDNA mixture (20 min, 37°C); GSSS cDNA was stored at -20°C before PCR. The PCR reactions were purified (Monarch PCR Cleanup Kit, New England Biolabs #T1030S) and amplification products visualized and quantified on ethidium bromide-stained gel and Bio-Rad Quantity One Analysis Software.

This locus contains two SNPs in linkage disequilibrium along the same haplotype: rs4841132 and a nearby SNP rs4841133. We took advantage of both variants to design primers for allele-specific RTPCR, permitting the use of two PCR amplicons that discriminate cleanly between major and minor cDNA. Both SNPs (rs4841132 and rs4841133) have G and A alleles, and both polymorphisms served as the most 3' base of their respective corresponding allele specific primers (where the 3'-ends are T for minor or C for major). Aliquots of the resulting cDNA were PCR amplified at high stringency annealing (66°C) with one primer in exon 1 (LOC157273_X1F; CAGCGAGCCATTTCAGCAGATTG) that annealed perfectly to both major and minor lncRNAs and rs4841132 primers that annealed to the rs4841132-A allele (CTCTCAGGTCACCAGCTGGAT) or the rs4841132-G allele (CCCTCTCAGGTCACCAGCTGGAC). To obtain the longer product priming at the rs4841133 site, we use allele-specific primers for rs4841132-A (GTAGACCACAAGGTGGAAATGGT) or rs4841132-G (GTAGACCACAAGGTGGAAATGGC). We alternatively quantified allele-specific cDNAs by using general primers followed by cloning and sequencing to enumerate the relative abundance of minor and major allele cDNA clone and observed consistent results (See Supplementary Text Section "Alternative quantification of allele-specific LOC157273 transcription using cDNA cloning" and **Supplementary Table 3**).

#### Human hepatocyte sources and culture

Cryopreserved primary human hepatocytes were obtained from commercial sources and included the products HMCPTS (ThermoFisher Scientific) and HUCPI6 (Lonza) (**Supplementary Table 1**). Cryopreserved vials were thawed partially at 37°C and transferred to 45 to 50 mL of pre-warmed (37°C) Cryopreserved Hepatocyte Recovery Medium (ThermoFisher Scientific #CM7000) with gentle mixing. Hepatocytes were collected by centrifugation at 110 x g for 6 min and resuspended in complete Williams E medium with GlutaMax (ThermoFisher Scientific #32551020), supplemented with dexamethasone, penicillin-streptomycin, insulin, selenium, transferrin, BSA and linolenic acid (ThermoFisher Scientific #CM4000). Cells for RNA analysis were plated in 2 mL of complete medium in 6-well collagen-coated plates (ThermoFisher Scientific #A1142801) at ~1 million cells per well. Between 6 and 16 hours later, loose cells were removed from the monolayers by gentle trituration, and the medium was replaced with warmed Williams E complete medium additionally containing 1X HepExtend (ThermoFisher Scientific #A2737501).

#### Screen of human hepatocyte donors to identify rs4841132 genotype

We genotyped rs4841132 in the human hepatocytes with a 2.9 kb LOC157273 amplicon from purified DNA. DNA was prepared from 500,000 to 1,000,000 human hepatocytes after partial thawing of a cryovial and recovery by centrifugation, followed by one wash with 1X PBS. DNA was purified from cells using the DNeasy Blood & Tissue Kit (Qiagen 69504) with a yield of 10-24 µg. We designed PCR primers to amplify a 2.9 kb *LOC157273* gene-body amplicon (gLOC157273_outerF2, 5'-AGCAAAAACTGATCCGGAAG-3'OH and gLOC157273_outerR2, 5' TTCAGCCTCCATGGTAAAGG-3'OH, **Supplementary Table 2)**. Based on human reference genome GRCh37/hg19, the 2904 bp gene-body amplicon [chr8:9,182,852-9,185,755] features 73 bp of exon 1, all 489 bp of intron 1, all 403 bp of exon 2, and 1939 bp of intron 2 parsed according to Ensembl transcript ENST00000520390.1. Genomic DNA (500 ng) was amplified in a 25 µL PCR reaction using

Expand™ Long Template PCR System and Buffer 1 (Sigma-Aldrich 11681834001), using an elongation temperature of 68°C and an annealing temperature of 60°C. After separation on a 1% agarose gel (1X TAE), bands were visualized after staining in the fluorescent dye SYBR-Gold on the Safe Imager™ 2.0 Blue-Light Transilluminato (ThermoFisher G6600) and excised with a razor blade. Uncloned DNA was gel-purified using Qiagen QIAquick Gel Extraction Kit and sequenced directly at Genewiz (South Plainfield, NJ) using our custom LOC157273 primers (seq4, seq6, seq7; see **Supplementary Table 9**).

#### Bioinformatic evidence that *LOC157273* is a candidate causal transcript

We retrieved the GRCH37/hg19 human genome reference assembly coordinates for all exons of ENCODE Consortium-catalogued long non-coding RNA genes, containing 13,562 genes and 23,105 transcripts (GENCODE v18, April 2013, www.gencodegenes.org) (3). All UCSC Genome Browser coordinates in the manuscript refer to the GRCh37/hg19 assembly of the human genome. We also retrieved the exon coordinates for all 81,673 mRNAs representing 20,318 protein-coding genes (GENCODE v18, April 2013) and intersected both sets of exons with all significant disease-associated GWAS SNPs listed in the NHGRI-EBI GWAS Catalog (http://www.ebi.ac.uk/gwas/docs/methods). Using the UCSC Genome Browser, we manually annotated each locus where a GWAS Catalog-listed SNP resided in a Gencode lncRNA exon. We excluded associations that fit at least one of the following conditions: (i) mapped inside a segmental duplication, copy number variant, or RepeatMasker-identified repetitive element; (ii) overlapped any exons or introns of any NCBI RefSeq or Gencode-catalogued protein-coding genes; (iii) resided less than 10 kb away from the nearest protein-coding gene promoter or in Gencode lncRNAs that had clear evidence of protein-coding capacity and were (at the time of our analysis) misannotated as lncRNAs in Gencode annotations; or (iv) were not exonic to any Genbank cDNAs or ESTs.

By comparing genome locations for exons from lncRNAs and protein-coding genes in GENCODE v18 with variants associated with complex traits and diseases in the NHGRI-EBI GWAS Catalog, we found 41,153 variant-published GWAS pairs. Manually annotating the top 71 pairs in the UCSC Genome Browser, we found the Chr8p23.1 PPP1R3B-LOC157273 locus to have the greatest amount of published GWAS evidence for disease associations from among all Gencode lncRNAs (**Supplementary Table 4**).

To interrogate the genomic structure of the LOC157273 transcript (NR_040039.1; ENST00000520390.1), we viewed the locus using the UCSC Genome Browser GRCh37/hg19 (Feb. 2009) human genome assembly (4). We manually annotated the locus using the RefSeq and GenBank cDNA tracks, as well as the Gencode Comprehensive Gene Annotation Set (V19), Gencode Pseudogenes and 2-way Pseudogenes (V19), GTEx RNA-Seq Gene Expression (53 Tissues, 570 donors) (5), and Human Body Map of lincRNAs and TUCP Transcript Tracks (lncRNA expression, 22 human tissues and cell lines). UCSC Genome Browser viewing mode was set to “Full” for all these tracks where applicable. The Human ESTs and Human mRNA tracks and the following ENCODE Integrated Regulation Tracks were used in the manual annotation: Layered H3K4Me1 (regulatory elements), Layered H3K4Me3 (near promoters), Layered H3K27Ac (active regulatory elements), DNaseI Clusters (125 cell types, ENCODE V3), Txn Factor ChIP (161 factors), and Txn Fac ChIP V2. We used HaploReg v4.1 (6) to explore the regulatory annotations of the haplotype on which rs4841132 resides. To link potential regulatory networks to the lncRNA transcripts arising from the LOC157273 locus, we interrogated the transcription start site, upstream region and gene body for tissue-specific DNaseI hypersensitivity sites, transcription factor binding sites, and the epigenetic marks H3K4me1, H3K4me3 and H3K27ac. We also interrogated the FANTOM Consortium HeliScope single-molecule RNA 5’ Cap Analysis of Gene Expression (CAGE) next-generation sequencing resource for the tissue-specificity of LOC157273 (7).

#### *LOC157273* conservation

We evaluated *LOC157273* conservation between human and mouse and performed reciprocal BLASTN and BLAT analysis of RepeatMasker-processed human *LOC157273* cDNAs, as described in our prior work (8). We also manually viewed the human *LOC157273* promoter and gene body in the UCSC Genome Browser (4), using the PhyloP and the 100-species MultiZ tracks. We assessed the evolutionary history of rs4841132 in ancient humans and nonhuman primates. We viewed each Denisovan and Neanderthal genome in the UCSC Genome Browser (at <100 bp resolution) to determine whether the nucleotide base at the rs4841132 position was G or A. We noted any discrepancies between the read and the GRCh37/hg19 human genome assembly. We used the UCSC Genome Browser's GWAS SNPs track clickthrough functionality to determine which allele of each SNP was the disease risk-associated allele, based on all published GWAS datasets reporting significant disease or trait associations of this SNP.

The PhyloP scores and MultiZ alignments of the human *LOC157273* promoter and gene body to 100 nonhuman species in the UCSC Genome Browser suggest that, while functional constraint has acted upon the *LOC157273* promoter, the exons as well as the introns of this lncRNA have been diverging at the neutral rate, consistent with the hypothesis that for lncRNAs, the act of their transcription may be functionally more important than the actual sequence of the transcript (**Supplementary Figure 4C-D**). Multispecies genome alignments in the UCSC Genome Browser show that most mammals - including all nonhuman primates, and except rodents and bats - have a G at the rs4841132 orthologous position, as did all of the ~ 25 Denisovan reads and one of two Neanderthal reads (**Supplementary Figure 4E**), suggesting that the rs4841132 A allele may have arisen during recent human evolution.

#### Alternative quantification of allele-specific *LOC157273* transcription using cDNA cloning

In Figure 5 in the main document, we use four different allele specific PCR primers (**Supplementary Table 3**) to address the question of the abundance of *LOC157273* transcripts from the major and minor copies in the TRL4079 heterozygote. To confirm the relative contribution of major and minor transcripts from this first approach, we took a parallel and alternative approach to all *LOC157273* cDNAs from the A/G heterozygote, followed by sequencing of plasmids to reveal major or minor. To this end, we primed synthesis of a *LOC157273* cDNA using a gene-specific strand-specific (GSSS) primer that anneals to both major and minor transcripts (LOC157273_revTprimer1; 5'-GGAAGACCACCAAAGAGATTATTC-3'-OH). In first strand synthesis, we annealed the revT1primer1 to 5 µg RNA from our A/G heterozygote (TRL4079) at 70°C in the presence of dNTPs and cooled on ice prior to cDNA synthesis. Together, the primers revT1primer1 (779) and LOC157273Fprimer1 (777; 5'-GACCCAGACATCATGGGAACTTC-3'-OH) amplify a 1.1 kb cDNA amplicon. After RNAse H treatment, the subsequent PCR reaction utilized the thermostable polymerase Platinum Blue (Invitrogen; 21.25 µL), 1.25 µL cDNA template, and 1.25 µL of each primer (10 µM each). Annealing was 53°C (30 sec), followed by 72°C (1 min 10 sec), 94°C (45 sec) for a total of 30 cycles; a final 10 min 72°C step was used. The resulting 1.12 kb band was visualized on an EtBr-agarose gel, excised with a razor blade, and purified with the Gel Extraction Kit (Qiagen). Purified DNA (15 ng) was ligated into pCR2.1TOPO in the TOPO TA Cloning kit (Invitrogen), and transformed into C2984 competent E coli cells (New England Biolab), selecting on LB-kanamycin plates with X-gal. White colonies were picked and miniprep DNA was digested with Eco R1 to identify plasmids with proper inserts.

Plasmids with *LOC157273* inserts were sequenced (Genewiz, South Plainfield, NJ), and aligned to GRCh37/hg19 using the UCSC Genome Browser, which revealed clones with the G or A substitutions at rs4841132 or rs4841133. This alternative approach (wholly different from the approach taken in Figure 5) involved sequencing 25 plasmid clones from the A/G heterozygote (nine clones Genewiz 30-25008756; additional sixteen clones Genewiz 30-26815569). Ten of 25 clones (40%) showed A at SNP rs4841132 (thus minor) and 15 out of 25 (60%) showed G. This skew towards cDNA clones with rs4841132 A supports the evidence for transcriptional crippling obtained using allele-specific PCR **(Figure 5)**.

#### Targeted resequencing of 5.6 kb of genomic DNA covering the gene body and upstream region of *LOC157273* in a single heterozygote hepatocyte donor

Two regions of the *LOC157273* totaling 5.6 kb were amplified by long-and-accurate PCR. In addition to the 2.9 kb gene-body region, we PCR-amplified the upstream region of *LOC157273* and acquired a 2.73 kb amplicon [chr8:9,180,153 to 9,182,914 on the reference genome GRCh37/hg19] including 346 bp of exon 1 extending 5' to -2418 bp relative to the transcriptional start site (TSS, capsite). The 2.9 kb "gene-body" and 2.73 kb “upstream” amplicons have a 65 bp overlap region in exon 1 (which spans 363 bp; chr8:9,182,561-9,182,923). Genomic DNA (500 ng) from the rs4841132 heterozygote (TRL4079) was amplified in a 25 µL PCR reaction using Expand™ Long Template PCR System and Buffer 1 (Sigma 11681834001), using an annealing temperature of 60°C and an elongation temperature of 68°C. The PCR primers were gLOC157273_promF1 (5’-ACTATGGCGAGCCAGAGATG-3’OH) and gLOC157273_promR1 (5’-GCTGAAATGGCTCGCTGAAC-3’OH). The amplicon includes the transcription-factor-binding region visible in the UCSC Genome Browser (GRCh37/hg19, chr8:9,182,104-9,182,560) located at -458 to -166 bp relative to the TSS. The plasmid used to clone fragments was pCR2.1TOPO (3931 bp; ThermoFisher, catalog 450641).

Pre-mix sequencing reactions containing 600-700 ng of purified plasmid DNA plus primer were submitted to GeneWiz (South Plainfield, NJ). The primers used for sequencing from plasmid-cloned amplicons are shown in **Supplementary Table 9**.

#### Plasmid-based overexpression of LOC157273 cDNA in primary human hepatocytes and hepatoma cells

To overexpress *LOC157273* as a full-length cDNA, we commercially obtained a mammalian expression construct (pTCN_LOC157273A, 8584 bp) featuring the CMV promoter driving a 3.4 kb cDNA insert (NR_040039.1), flanked by an upstream BamH1 site and a downstream Xba1 site (transOMIC Technologies; Huntsville, AL). The construct harbors the A minor alleles at SNP rs4841132 and SNP rs4841133. The 8.6 kilobase plasmid was verified by partial sequencing before use. We created the 5.2 kb empty vector (pTCN; TransOmic pTCN Expression Vector) by removing the 3.4 kb BamH1-Xba1insert.

Both primary human hepatocytes (Hu8200 donor) and human Huh-7 hepatoma cells (9) were targeted for the overexpression plasmid described above. In the case of the hepatoma line, cells were transfected with pTCN empty vector or pTCN LOC157273A (both carrying the neomycin/kanamycin cassette); 500 µg G-418 (ThermoFisher Scientific #10131-035) was used to select stable tranfectants of the empty vector and 8.6 kb expression plasmid based on kill-curve performance antibiotic concentration for clonal selection. For the production of stable clones, transfected cells were allowed to recover for 48 hr before addition of G-418. Subsequently, cells were passaged at 1:10 in Dulbecco's Modified Eagle Medium (DMEM) containing 10% fetal bovine serum and 500 µg G-418/mL, and G-418-containing media replaced every 3 to 4 days until resistant clones appeared at 14 days. G-418 resistant colonies were trypsinized and cultured first in 24-well plates before expansion to 100 mm dishes for the preparation of RNA. RNA was prepared using RNeasy Kits (Qiagen #74104). One empty vector clone was maintained as well as eight *LOC157273* overexpression clones. Total RNA was transcribed into cDNA using oligo(dT) priming, and TaqMAN real-time RT-PCR done using primers for *GAPDH*, *PPP1R3B* and *LOC157273*. Values for $\Delta C_T$ were computed relating *PPP1R3B* or *LOC157273* to *GAPDH* expression.

Expression of the LOC157273 lncRNA in primary human hepatocytes is low compared to most mRNAs; we estimated ~50 copies per cell in hepatocytes homozygous for the rs4841132 G allele. We overexpressed LOC157273 using G-418-resistant clones harboring either empty vector (pTCN) or pTCN_LOC157273A (where the CMV promoter drives expression of the 3.4 kb LOC157273 cDNA). We did not observe consistent responses in *PPP1R3B* expression compared to an empty vector in either human hepatocytes (**Supplementary Figure 6A**) or in the human hepatoma cell line (Huh7) (**Supplementary Figure 6B**).

#### Supplementary Figures

##### Supplementary Figure 1: Screen of the LOC157273 region from 16 human hepatocyte donors to identify rs4841132 genotype

**Supplementary Figure 1a:** Nucleotide assignments of seven published SNPs and one indel defining the major and minor ***LOC157273*** haplotype of donor TRL4079.

DNA purified from hepatocyte donor TRL4079 was amplified using Expand Long Template polymerase (Sigma-Aldrich Roche 11681834001) targeting a genomic amplicon of 2.9 kb (*hg19* chr8:9,182,852-9,185,756). The gel-purified DNA was cloned into pCR2.1TOPO. Clones representing the major (clones 2, 6, 7) and minor (clones 3, 4, 5) alleles were sequenced using five primers (including M13forward and M13reverse), and base choices assigned for seven SNPs and the indel rs111675407 in the first intron. Gene-specific primers are *seq2* (forward; 5'-CACGTGGTACTGGCACTCCGTTTG-3'OH), *seq8* (forward; 5'-GGTCTTTCCTCTATGCCAGGATTG-3'OH), *seq5* (reverse; 5'-TCCATGGTAAAGGCTGGGACTTG-3'OH) and *seq6* (forward; 5'-CTTGACACCATCTTCAGTAATTTC-3'OH).

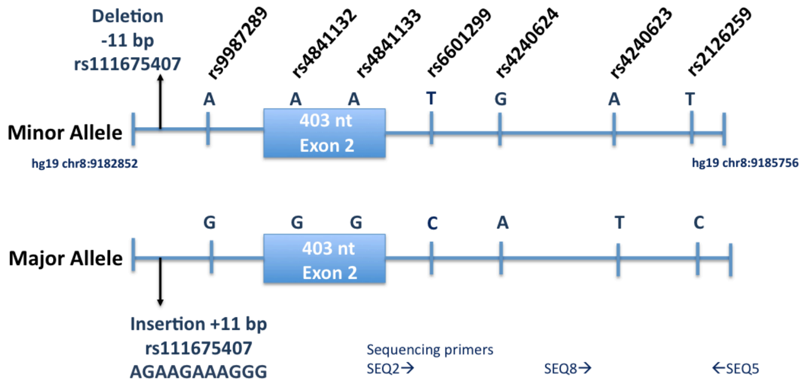


##### Supplementary Figure 1b: DNA resequencing of major and minor alleles of the LOC157273 shows heterozygosity at SNP rs2169387 (-1170 relative to the transcriptional start site), a SNP associated with serum total cholesterol.

DNA from hepatocyte donor TRL4079 was amplified using a pair of primers designed to amplify a 2.76 kb amplicon (reference genome hg19 chr8:9,180,153-9,182,914) in the upstream region; gel-purified DNA was cloned into the plasmid pCR2.1TOPO. Two plasmid clones (#7 and #10) were sequenced (one each representing minor and major versions) using sequencing primers: LOC157273_upstream_F1 (5'-GTGCCTGGCCATAGTCATCTC-3-OH) and LOC157273_upstream_R1 (5'-ATGTTTTCCCCCGGTTCAGAG-3'OH). Sequence files were aligned to human reference genome hg19 using BLAT; the vertical red bars indicate that the BLATted file differs from than the reference genome (which happens to be a minor allele version) at four positions in the 300 bp region shown. The SNP variant rs2169387 (G or A; chr8:9,181,395) is located in a predicted liver and muscle enhancer region; the minor allele is associated with elevated total cholesterol (10). Clone 10 has G at -1166 (major); clone 7 has A (minor) at this position (-1166).

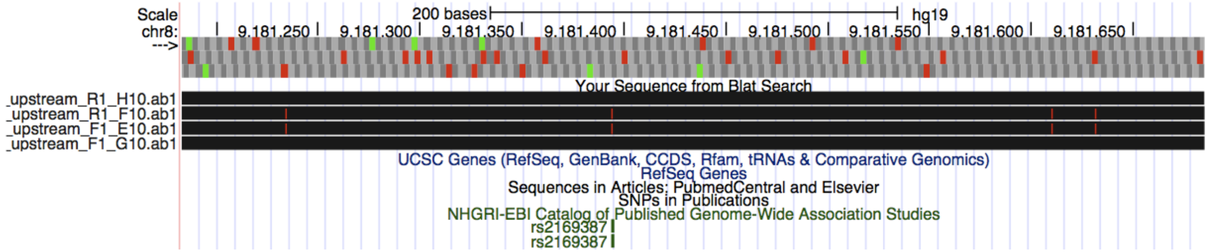


**Supplementary Figure 1c: Sanger sequence reads for a LOC157273 heterozygote hepatocyte donor show half-height A/G peaks at both SNP rs4841132 and rs4841133.**

**Panel A:** The region of the chromatogram surrounding rs4841132 is shown, where two half-height peaks (green A, black G) at position 83 are visible.

**Panel B**: The region surrounding rs4841133 is shown; note the two half-height peaks at position 151 (rs4841133).

A

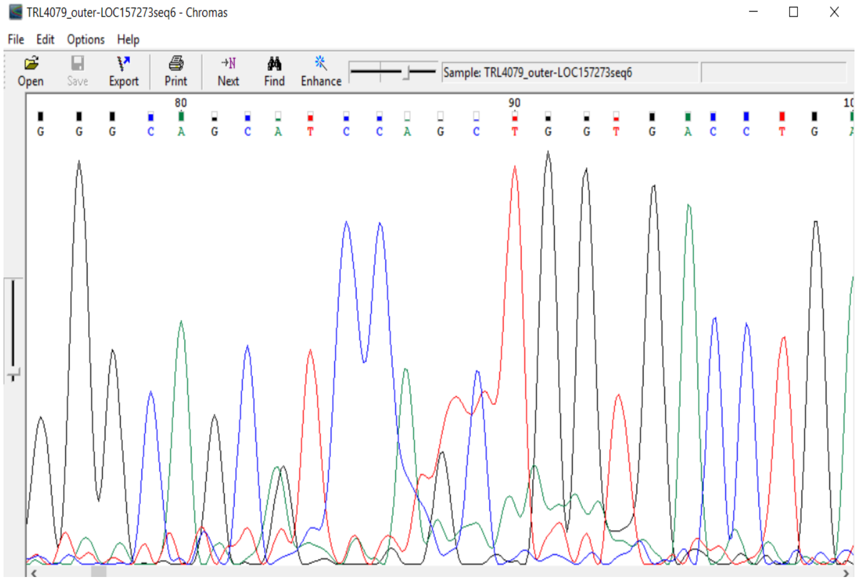


B

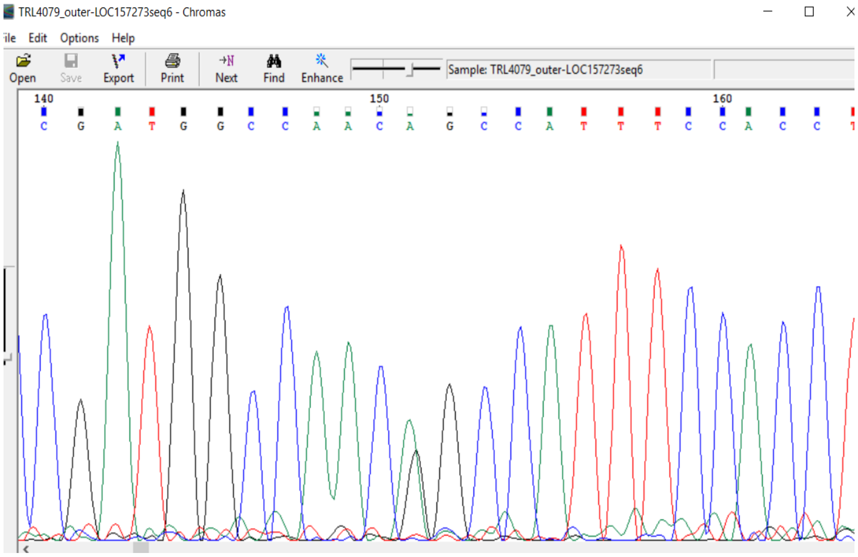

**Supplementary Figure 2: Genetic associations with fasting glucose and fasting insulin at the chromosome 8 PPP1R3B-LOC157273 locus in the MAGIC, AAGILE and CHARGE consortia.**

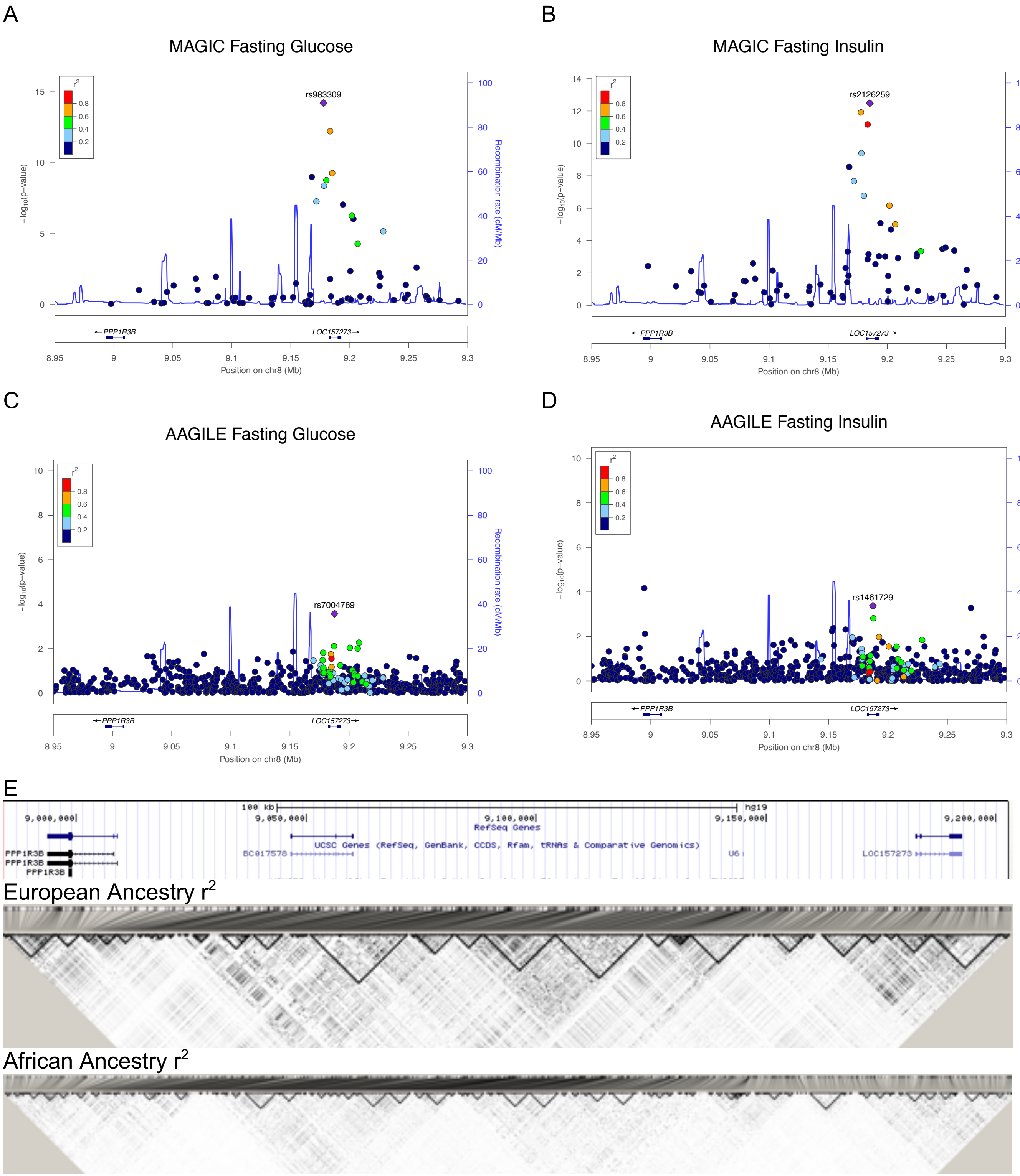

**Panel A:** Regional association plot of the chromosome 8 *PPP1R3B-LOC157273* locus associated with **fasting glucose in European** ancestry individuals in MAGIC. Data are from Liu et al AJHG 2016. Statistical significance of the association of each SNP (dots) is shown on the Y-axis (-log(*P*-value)) and chromosome 8 genomic position on the X-axis. The purple diamond (rs983309, $r^2$ with rs4841132 = 0.74) is the most significantly-associated SNP ($p=6.29x10^{-15}$) in the region, with the color of each dot indicating LD ($r^2$) with the most significant SNP based on 1000 Genomes Reference data. The blue line represents the recombination rate. rs4841132 lies in the body of LOC157273.

**Panel B**: Regional association plot of the chromosome 8 *PPP1R3B-LOC157273* locus associated with **fasting insulin in European** ancestry individuals in MAGIC. The most significantly-associated SNP (rs2126259, $p=3.13x10^{-13}$) is highly correlated with rs4841132, red diamond, $r^2=0.83$). Other details are the same as in Panel A.

**Panel C**: Regional association plot of the chromosome 8 *PPP1R3B-LOC157273* locus associated with **fasting glucose in African-American** ancestry individuals in AAGILE. The purple diamond (rs7004769, $r^2$ with rs4841132=0.44 in African ancestries) is the most significantly-associated SNP ($p=2.69x10^{-4}$) in the region, with the color of each dot indicating LD ($r^2$) with the most significant SNP based on Africans in the 1000 Genomes Reference data. Other details are the same as in Panel A.

**Panel D**: Regional association plot of the chromosome 8 *PPP1R3B-LOC157273* locus associated **with fasting insulin in African-American** ancestry individuals in AAGILE. The purple diamond (rs1461729, $r^2$ with rs4841132 = 0.96) is the most significantly-associated SNP ($p=4.2x10^{-4}$) in the region, with the color of each dot indicating LD ($r^2$) with the most significant SNP based on Africans in the 1000 Genomes Reference data. Other details are the same as in Panel A.

**Panel E.** Regional linkage disequilibrium across the chromosome 8 *PPP1R3B-LOC157273* locus. The top panel shows the UCSC gene track, with *LOC157273* on the right and *PPP1R3B* on the left, the middle panel shows European ancestry LD (HapMap2 CEU) and the bottom, African ancestry (HapMap2 YRI). As expected, LD across the region is more fragmented in YRI than in CEU.

#### Supplementary Figure 3: DNAseI Hypersensitivity, Transcription Factor Binding, and Expression profile of *LOC157273*

**Supplementary Figure 3A**: DNAseI Hypersensitivity Clusters and Transcription Factor Binding in the promotor region of *LOC157273*.

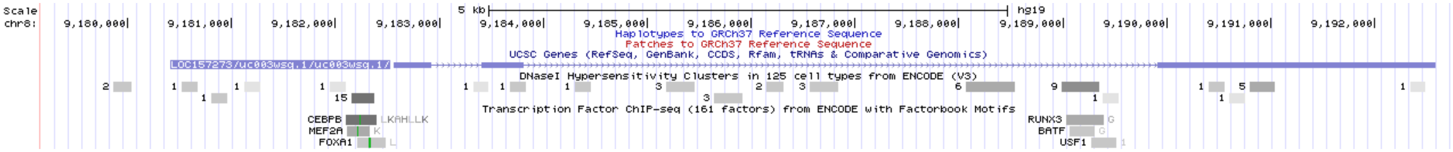


**Supplementary Figure 3B**: Mapped transcription start sites (TSS) and their usage in human and mouse primary cells, cell lines and tissues from FANTOM5 (http://fantom.gsc.riken.jp/5/). LOC157273 expression across the human body was queried with single molecule sequencing. The figure shows the 5'-end of the mapped CAGE reads counted at a single base pair resolution (CTSS, CAGE tag starting sites) on the genomic coordinates, which represent TSS activities in the sample. Identified peaks from the "CAGE Phase1 CTSS human tracks pooled, filtered with 3 or more tags per library and rle normalized" track shows that the strongest DNase I hypersensitive site lies at the conserved promoter of *LOC157273*. lncRNA *LOC157273* is almost exclusively expressed in human hepatocytes from several adult donors (three are adult human hepatocytes and one is total pooled adult liver.) Moreover, *LOC157273* is not appreciably detectable in nearly any other of the 1,000 human organs, tissues, cell types, or cell lines available in the CAGE human promoterome resource.

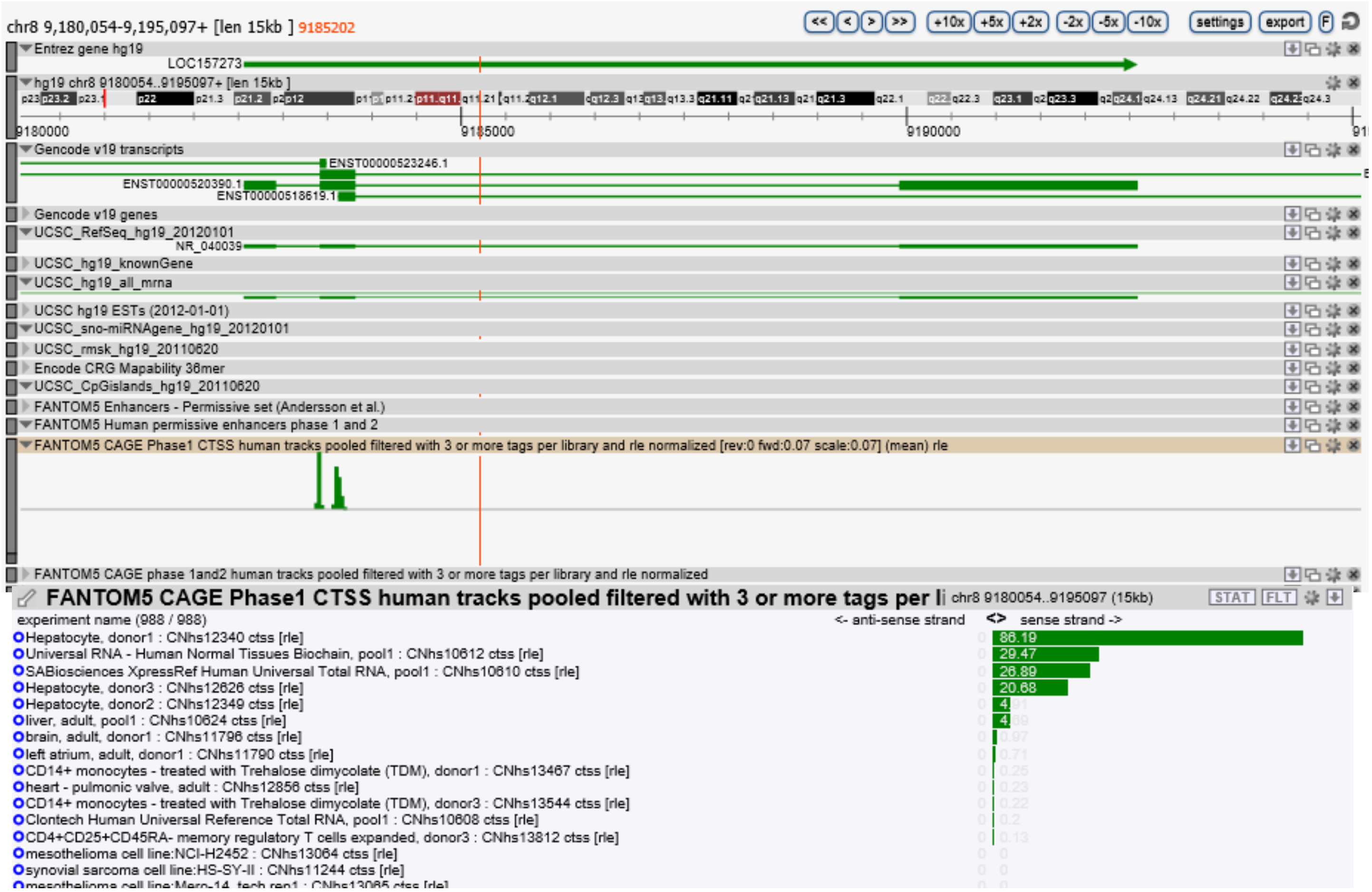

**Supplementary Figure 3C**: Gene Expression in 53 tissues from GTEx RNA-seq of 8555 samples. In the NIH-sponsored The Genotype-Tissue Expression (GTEx) Project, 53 tissues were surveyed for expression of the lncRNA *LOC157273*. The only significant positive tissue was liver (light brown peak), median RPKM = 3.040. By comparison, the expression of GAPDH in liver (median) is 571.0; for PPP1R3B in liver (median) is 42.29. Assuming an absolute transcript abundance for GAPDH of 1250 copies per cell, the *LOC157273* lncRNA would be estimated at 6.7 copies per cell.

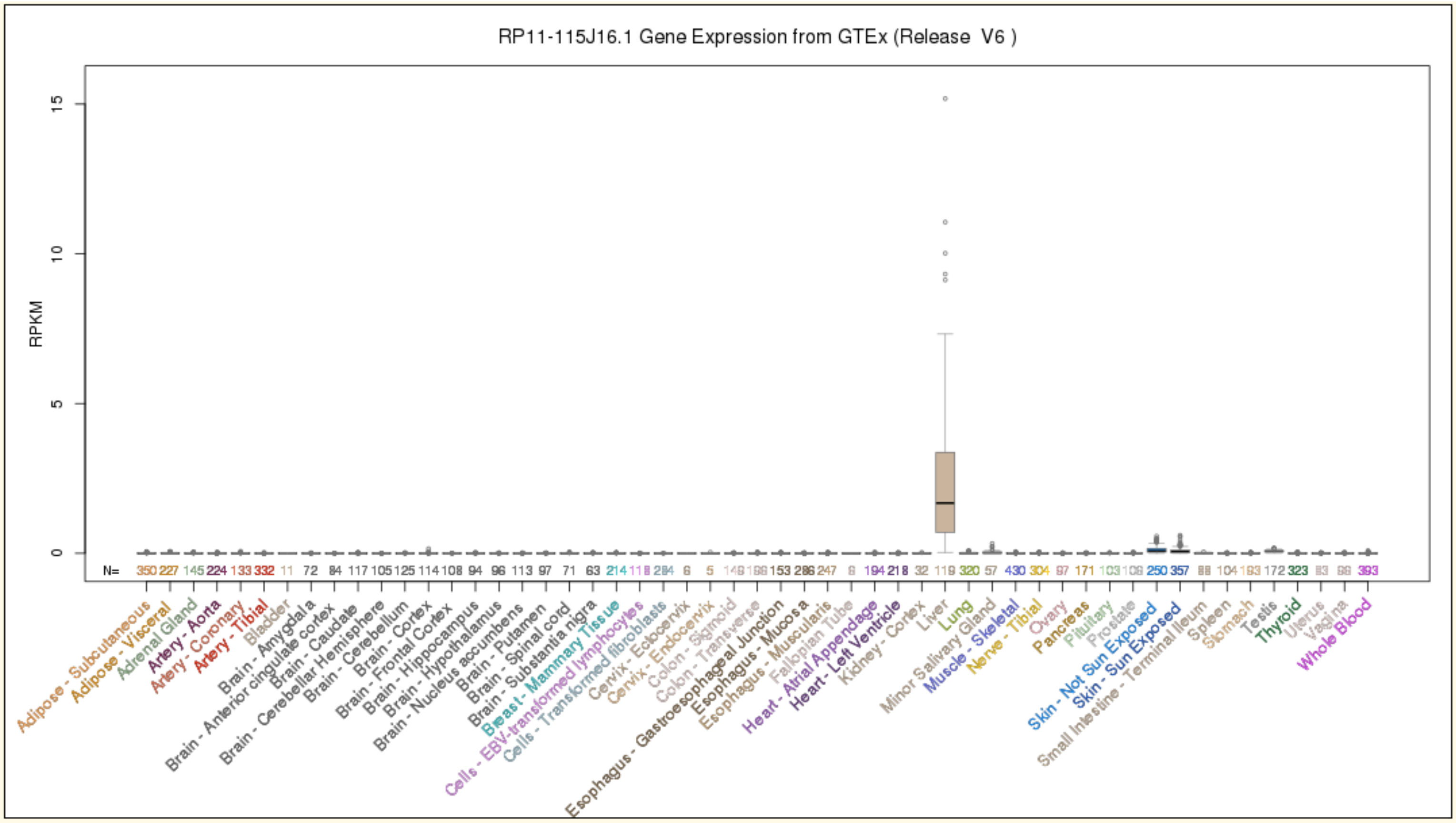

**Supplementary Figure 4: Comparative genomics of *LOC157273* in modern and ancient humans, nonhuman primates, and mouse.**

**Supplementary Figure 4A**: UCSC BLAT (no RepeatMasker) search of human hg19 *LOC157273* query versus mouse mm9 shows that the top hit (score 44, first line) is extremely short. (With RepeatMasker, human *LOC157273* does not have any BLAT hits in mouse.)

**Mouse BLAT Results**

**BLAT Search Results**

| ACTIONS | QUERY | SCORE | START | END | QSIZE | IDENTITY | CHRO | STRAND | START | END | SPAN |
|---|---|---|---|---|---|---|---|---|---|---|---|
| browser details | YourSeq | 44 | 3377 | 3431 | 3442 | 91.0% | 7 | + | 80161087 | 80161142 | 56 |
| browser details | YourSeq | 27 | 3402 | 3434 | 3442 | 87.1% | 12 | - | 82152395 | 82152426 | 32 |
| browser details | YourSeq | 25 | 2616 | 2644 | 3442 | 85.8% | 4 | + | 142312285 | 142312312 | 28 |
| browser details | YourSeq | 22 | 3231 | 3256 | 3442 | 92.4% | 8 | - | 105881361 | 105881386 | 26 |
| browser details | YourSeq | 21 | 3382 | 3412 | 3442 | 83.9% | 10 | - | 66799371 | 66799401 | 31 |

**Supplementary Figure 4B**: Mouse Ppp1r3b is on chr 7, not chr 8. Since the *LOC157273* BLAT hit is on mouse chr 8, the mouse *LOC157273* putative homolog is not an ortholog, as it aligns to a non-orthologous mouse region relative to human LOC157273-PPP1R3B.

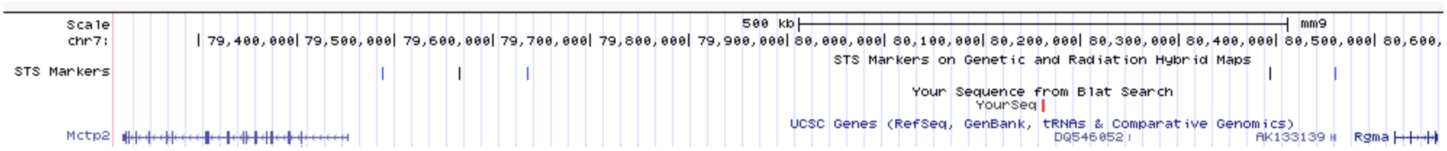


**Supplementary Figure 4C**: UCSC Browser PhyloP and MultiZ analysis shows that, although human *LOC157273* lncRNA has a conserved promoter, its exons are not well-conserved.

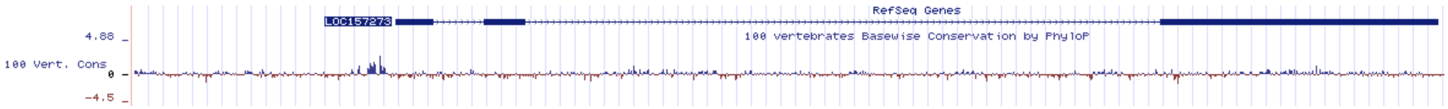

**Supplementary Figure 4D**: MultiZ Alignments of 100 vertebrates shows the extent of conservation across species for rs4841132.

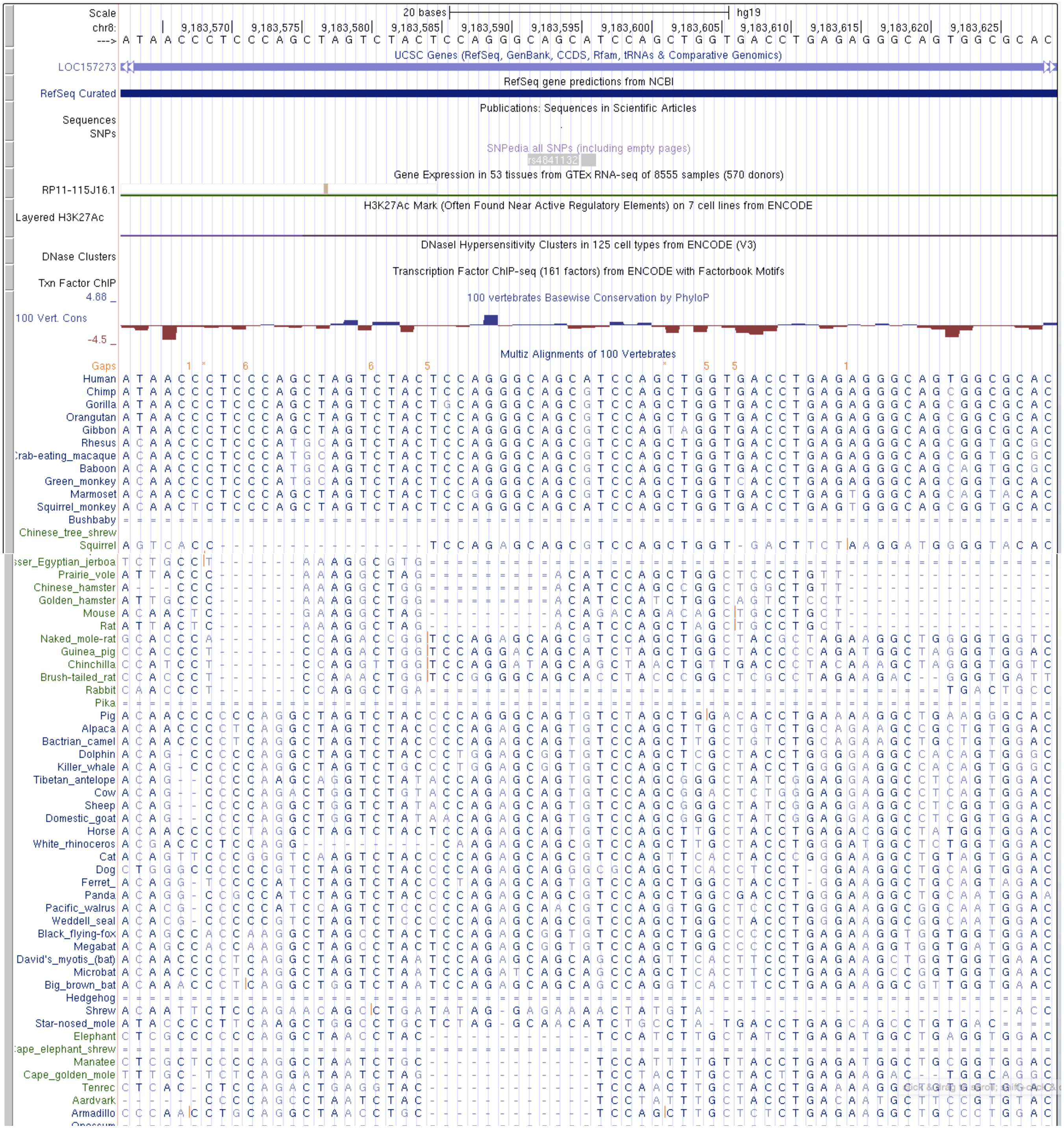

**Supplementary Figure 4E:** Sequence conservation around the risk allele rs4841132A occurs across humans, including Denisovans. The GWAS rs4841132A risk allele is not observed in Denisovans.

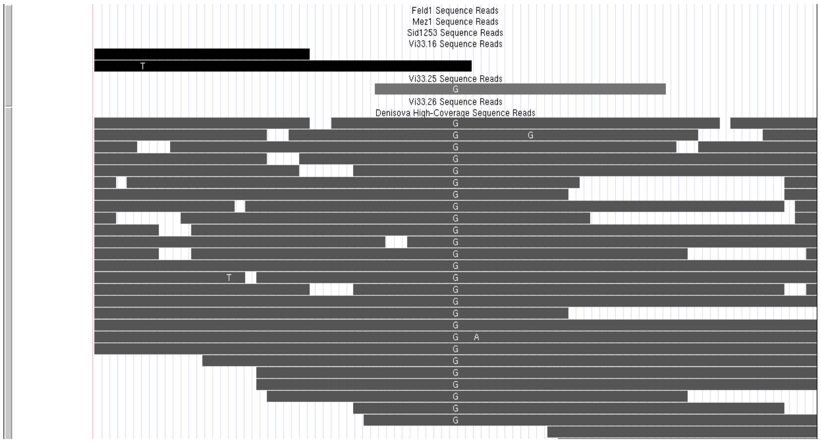

**Supplementary Figure 5: Glycogen content in human hepatocytes (Hu8200 donor homozygote G/G for rs4841132) in siRNA knockdown vs siRNA control conditions.**

Glycogen content was measured by incubating whole cell lysates with Aspergillus niger amyloglucosidase (See Supplementary Methods). The glucose concentration, estimated using a standard curve, is shown for Hu8200 primary hepatocytes treated with siRNA targeting *LOC157273* (siRNA11, siRNA15) or control siRNA (Mock, Scramble). Data are shown for three replicate experiments.

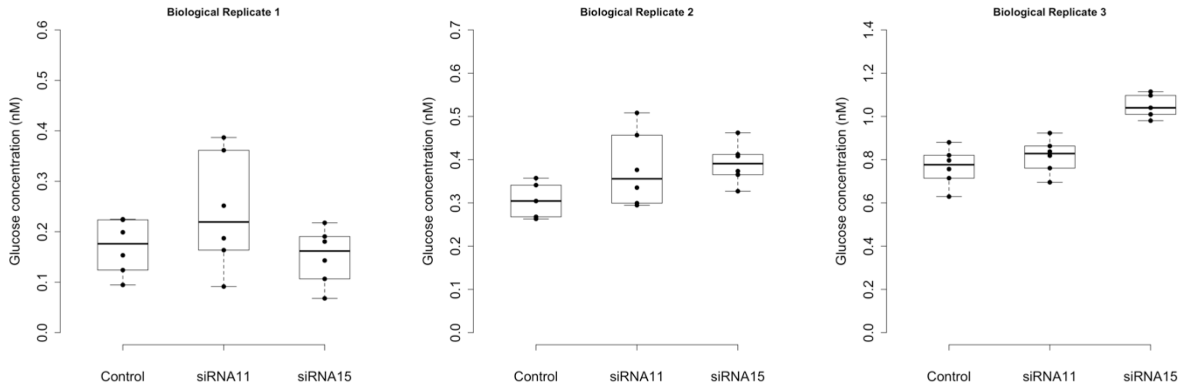

**Supplementary Figure 6: Fold-change in *LOC157273* and *PPP1R3B* expression after transfection with plasmid to overexpress *LOC157273***

Cells was transfected with either empty vector (pTCN) or an expression vector (pTCN_LOC157273A) harboring the 3.4 kb *LOC157273* cDNA. Total RNA was transcribed into cDNA using oligo(dT) priming, and TaqMAN real-time RT-PCR done using primers for GAPDH, PPP1R3B and *LOC157273*. Values for $\Delta C_T$ were computed relating *PPP1R3B* or *LOC157273* to GAPDH expression.

**Panel A:** Response in *PPP1R3B* gene expression in human hepatocytes (donor Hu8200) transfected with pLOC plasmid compared to empty vector. Mean "$\Delta\Delta C_T$ of *PPP1R3B*" is 1.06, median is 0.995, interquartile range is (0.96, 1.17). Spearman's rank correlation coefficient between "$\Delta\Delta C_T$ of LOC157273" and "$\Delta\Delta C_T$ of *PPP1R3B*" is 0.08 (P=0.91).

**Panel B:** Response in *PPP1R3B* gene expression in human hepatoma cell line (Huh-7) transfected with pLOC plasmid compared to empty vector. Mean "$\Delta\Delta C_T$ of *PPP1R3B*" is 2.24, median is 1.6, interquartile range is (1.08, 2.0). Spearman's rank correlation coefficient between "$\Delta\Delta C_T$ of LOC157273" and "$\Delta\Delta C_T$ of *PPP1R3B*" is -0.42 (P=0.41).

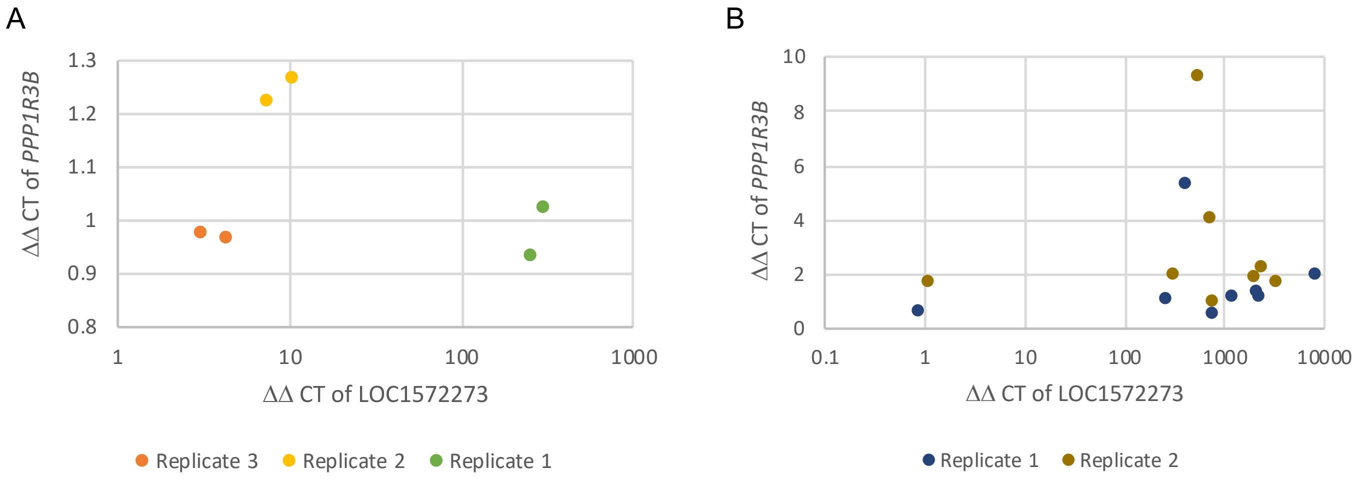

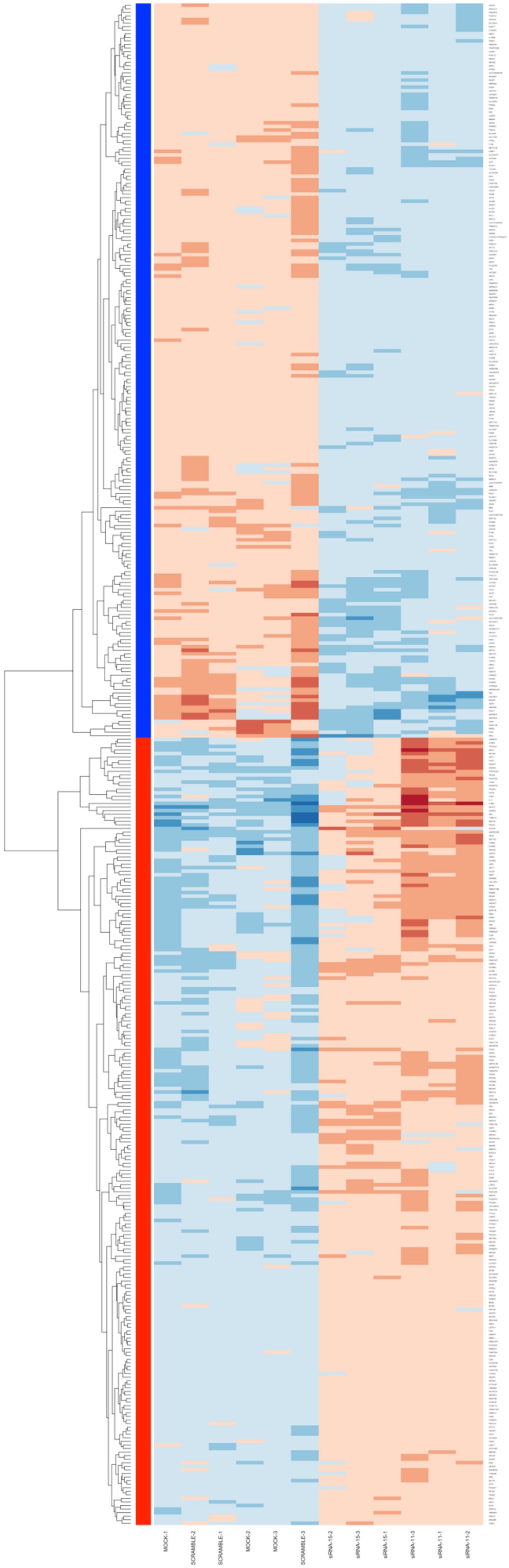


**Supplementary Figure 7. Cluster analysis showing clustered genes with similar normalized transcript counts in siRNA11, siRNA15 and control conditions from the transcriptome-wide differential expression analysis.**

The dendrogram shows relative similarity among genes. The analysis was restricted to differentially expressed genes with a relative Fold Change > 1.34 (first column on left after dendrogram, red row markers indicating higher counts compared to the mean count of each gene), or Fold Change < 0.74 (blue row markers indicating lower counts compared to the mean count of each gene), and $P_{adj} < 0.001$. Each row represents one gene and each column represents one of three replicates treated with siRNA targeting *LOC157273* (siRNA11, siRNA15) or control siRNA (Mock, Scramble).

**Supplementary Figure 8: Insulin re-stimulation of primary human hepatocytes stimulates glycogen deposition in hepatocytes from additional heterozygous (G/G) donors.**

**Panel A** shows the response in TRL4113 primary hepatocytes, with median basal glucose concentration of 0.14 nM and a no increase in glycogen content after insulin re-stimulation (P=0.65).

**Panel B** shows the response in TRL4012 primary hepatocytes, with median basal glucose concentration of 0.069 and no observable increase in glycogen content after insulin re-stimulation (P=1).

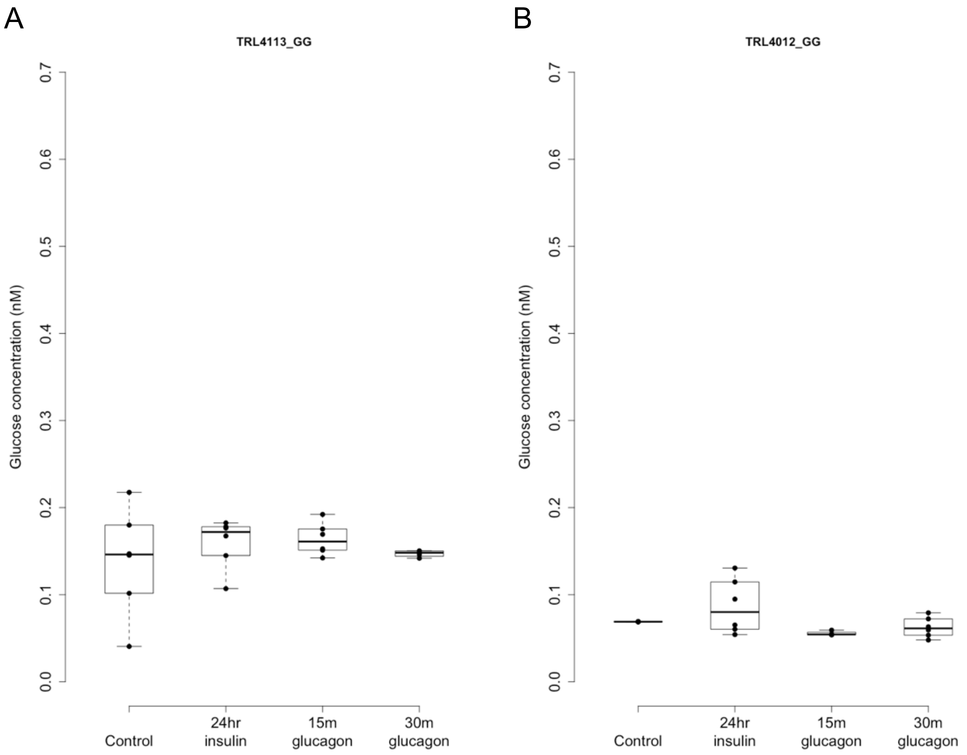

**Supplementary Figure 9: Physiological Model of *LOC157273* action on *PPP1R3B*, hepatic glycogen flux and peripheral glucose and insulin levels**

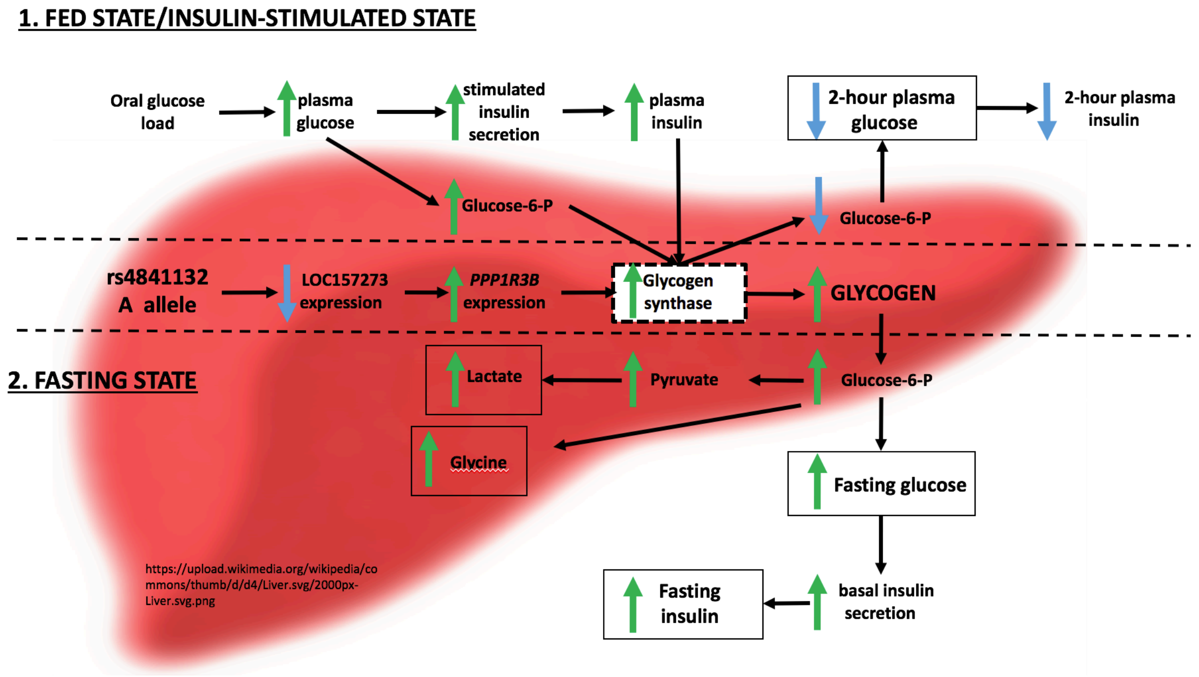

### Supplementary Tables

**Supplementary Table 1: Primary Human Hepatocyte Donor Characteristics**

| Vendor | LOT# | Ethnicity | Gender | rs4841132 Genotype | Age (years) | BMI (kg/m$^2$) | Cause of death |
|---|---|---|---|---|---|---|---|
| Triangle Research Labs (TRL) * | TRL4152 | Caucasian | Male | G/G | 18 | 24.3 | pain killer overdose |
| TRL | TRL4111A | Caucasian | Male | G/G | 27 | 31.6 | heroin overdose |
| TRL | TRL4107 | Caucasian | Female | G/G | 33 | 25.0 | drug overdose |
| TRL | TRL4094 | Caucasian | Male | G/G | 15 | 23.0 | head trauma |
| TRL | TRL4098 | Caucasian | Male | G/G | 24 | 22.8 | anoxia |
| TRL | TRL4113 | Caucasian | Male | G/G | 29 | 27.5 | anoxia |
| TRL | TRL4119A | African American | Female | G/G | 30 | 33.2 | cardiac arrest |
| TRL | TRL4000 | African American | Male | G/G | 39 | 19.0 | cardiac arrest |
| TRL | TRL4108 | African American | Female | G/G | 42 | 35.0 | stroke |
| TRL | TRL4133 | Caucasian | Female | G/G | 45 | 20.8 | anoxia |
| TRL | TRL4105A | Caucasian | Male | G/G | 45 | 24.2 | anoxia |
| TRL | TRL4055A | Caucasian | Female | G/G | 54 | 26.2 | stroke |
| TRL | TRL4012 | Caucasian | Male | G/G | 54 | 29.0 | stroke |
| TRL | TRL4056B | African American | Male | G/G | 55 | 28.9 | stroke |
| TRL | TRL4079 | Caucasian | Male | A/G | 24 | 31.5 | stroke |
| LifeTech | Hu8200 | African American | Male | G/G | 52 | 17 | anoxia |

* Triangle Research Labs is now part of Lonza.

**Supplementary Table 2. TaqMan primers chosen for two PPP1R3B mRNA known isoforms and a single set for LOC157273**

| Amplicon (LifeTech catalog #) | Intron spanned | Sequence in amplicon upstream of intron* | Intron sequence | Sequence in amplicon downstream of intron* |
|---|---|---|---|---|
| LOC157273 [Hs01371902_g1] | LOC157273 Intron 1 (490 nt) | ctccaatctgctgaaatggctcg ctgaactggaagg (36 nt) | /GTGGGC…TTGCAG (490 nt) | gcctcacccttggagatcagcac gtgctgcaggcaggggat (41 nt) |
| PPP1R3B "general" [custom order lot P160826-004-B12] | PPP1R3B Intron 1B (8895 nt) | tgcgggctctccgatgcggcga gcgagctggggagggggcttc tccgcggcccaaaag (58 nt) | /GTGGGTGCGGCGC GAGGCCAAGGA … TCCCTTTCCACGGT TCTAG (8895 nt) | cctgttcatctagccccatgatgg ctgtggacatcgagtacagatac aactgcatgg (57 nt) |
| PPP1R3B-HS (hepatocyte-specific) | PPP1R3B Intron 1A (9657 nt) | cgggttcagctcataagtcccg agggcaga (30 nt) | /GTACGTAGCTCC … TCCACGGTTCTAG/ (9657 nt) | cctgttcatctagccccatgatgg ctgtggacatcgagtacagatac aactgcatgg (57 nt) |

* The sense strand is shown in the table. The precise amplicons for each primer and probe-set were verified experimentally by sequencing plasmid hardcopies of qRT-PCR reaction products after cycle 36.

**Supplementary Table 3: Primers used in allelic-imbalance of LOC157273 transcription in a heterozygous hepatocyte donor**

| Name | Purpose | Region in *LOC157273* gene | Sequence of primer or siRNA genomic target |
|---|---|---|---|
| LOC157273_ revTprimer1 | Priming cDNA for cloning | RefSeq exon 3 (chr8:9,190,282-9,190,305) antisense to lncRNA | GGAAGACCACCAAAGAGATTATTC |
| gLOC157273_ seqR9 | Priming cDNA for allele-specific transcription | RefSeq exon 2 (chr8:9,183,692-9,183,715) antisense to lncRNA | AACCTCCATGCAGTCGGGATGAAG |
| LOC157273_ X1F | PCR primer for allele-specific transcription analysis | RefSeq exon 1 (chr8:9,182,898-9,182,920) | CAGCGAGCCATTTCAGCAGATTG |
| 32A* | PCR primer for allele-specific transcription analysis | Exon 2 | CTCTCAGGTCACCAGCTGGAT |
| 32G* | PCR primer for allele-specific transcription analysis | Exon 2 | CCCTCTCAGGTCACCAGCTGGAC |
| 33A* | PCR primer for allele-specific transcription analysis | Exon 2 | GTAGACCACAAGGTGGAAATGGT |
| 33G* | PCR primer for allele-specific transcription analysis | Exon 2 | GTAGACCACAAGGTGGAAATGGC |

*`Each of the four starred (*) primers are reverse primers in exon 2, chosen to work with the X1F forward primer in exon 1. The two primers at the top of the table are reverse primers used to enrich synthesis of *LOC157273* cDNA in a gene-specific strand-specific (GSSS) protocol.

**Supplementary Table 4: Genomic locations after intersection of GENCODE v18 exons with GWAS SNPs**

| Chromosome | Exon start position (b37) | Exon end position (b37) |
|---|---|---|
| chr1 | 1079198 | 1079199 |
| chr1 | 25296743 | 25296744 |
| chr1 | 25297184 | 25297185 |
| chr1 | 53581670 | 53581671 |
| chr1 | 101540999 | 101541000 |
| chr1 | 110439480 | 110439481 |
| chr1 | 164739171 | 164739172 |
| chr1 | 165180089 | 165180090 |
| chr10 | 8093034 | 8093035 |
| chr10 | 86953327 | 86953328 |
| chr10 | 101287764 | 101287765 |
| chr11 | 203788 | 203789 |
| chr11 | 2169774 | 2169775 |
| chr11 | 2691471 | 2691472 |
| chr11 | 90586900 | 90586901 |
| chr11 | 126030615 | 126030616 |
| chr12 | 58162739 | 58162740 |
| chr12 | 121438844 | 121438845 |
| chr13 | 75128114 | 75128115 |
| chr13 | 114622597 | 114622598 |
| chr16 | 52586341 | 52586342 |
| chr16 | 68600351 | 68600352 |
| chr17 | 46123004 | 46123005 |
| chr17 | 46123932 | 46123933 |
| chr17 | 70594629 | 70594630 |
| chr17 | 73872948 | 73872949 |
| chr18 | 905125 | 905126 |
| chr18 | 53855525 | 53855526 |
| chr18 | 56752054 | 56752055 |
| chr19 | 10397403 | 10397404 |
| chr19 | 12146370 | 12146371 |
| chr19 | 15724203 | 15724204 |
| chr19 | 21666210 | 21666211 |
| chr2 | 8036260 | 8036261 |
| chr2 | 45188353 | 45188354 |
| chr2 | 74208362 | 74208363 |
| chr2 | 177042633 | 177042634 |
| chr2 | 219840891 | 219840892 |
| chr20 | 23612737 | 23612738 |
| chr20 | 58896715 | 58896716 |
| chr21 | 29475196 | 29475197 |
| chr3 | 14530721 | 14530722 |
| chr3 | 169497585 | 169497586 |
| chr3 | 186570453 | 186570454 |
| chr3 | 186573705 | 186573706 |
| chr4 | 111718067 | 111718068 |
| chr4 | 174515682 | 174515683 |
| chr4 | 187207381 | 187207382 |

| | | |
|---|---|---|
| chr5 | 9623622 | 9623623 |
| chr5 | 134572229 | 134572230 |
| chr5 | 137194762 | 137194763 |
| chr5 | 158759900 | 158759901 |
| chr6 | 19803768 | 19803769 |
| chr6 | 29943067 | 29943068 |
| chr6 | 30782002 | 30782003 |
| chr6 | 32358513 | 32358514 |
| chr6 | 32628428 | 32628429 |
| chr6 | 134158800 | 134158801 |
| chr8 | 9183596 | 9183597 |
| chr8 | 10334781 | 10334782 |
| chr8 | 22400989 | 22400990 |
| chr8 | 23082971 | 23082972 |
| chr8 | 82445224 | 82445225 |
| chr8 | 123706155 | 123706156 |
| chr8 | 143761931 | 143761932 |
| chr9 | 22029547 | 22029548 |
| chr9 | 136131188 | 136131189 |
| chr9 | 136131322 | 136131323 |
| chr9 | 136131415 | 136131416 |
| chr9 | 136132908 | 136132909 |
| chr9 | 136478355 | 136478356 |

**The following Supplementary tables are in Supplement.xlsx**

**Supplementary Table 5. Haploreg (v4) query of rs4841132.**

**Supplementary Table 6: Genes with Padj<0.01 from gene expression differential expression analysis of the whole transcriptome comparing siRNA-11 and siRNA-15 to control.**

**Supplementary Table 7: Enriched Reactome pathways from the genes showing increased expression in siRNA11 and siRNA15 vs control.**

**Supplementary Table 7A**: Enriched Reactome pathways from the genes with differential expression P < 0.01, organized by hierarchical clustering of gene counts into two groups: genes showing increased counts in siRNA11 and siRNA15 conditions compared to control (of the 512 tested, 511 had Entrez gene ids and were used as input) and genes showing decreased counts in siRNA11 and siRNA15 conditions compared to control (of 441 genes testes, 440 had Entrez gene IDs).

**Supplementary Table 7B**: Enriched Reactome pathways from the clusters showing increased or decreased gene expression in siRNA11 and siRNA15 conditions, after hierarchical clustering of genes showing effects and significance as large as *PPP1R3B*: absolute value of FC >= 1.34 and $P_{adj}$<0.001.

**Supplementary Table 8: Reactome pathways showing enrichment in differential expression from Gene Set Enrichment Analysis.**

**Supplementary Table 9: Primers used to re-sequence cloned *LOC157273* genomic amplicons**

| Primer name | Genome location (hg19) | Primer sequence |
|---|---|---|
| seq2 | chr8:9,184,472-9,184,495 | CACGTGGTACTGGCACTCCGTTTG |
| seq3 | chr8:9,185,151-9,185,173 | AACTGAGCCTGAGCTGTCTAGAG |
| seq4 | chr8:9,182,858-9,182,879 | AACTGATCCGGAAGACTTGTTG |
| seq5 | chr8:9,185,725-9,185,747 | TCCATGGTAAAGGCTGGGACTTG (reverse primer) |
| seq6 | chr8:9,183,497-9,183,520 | CTTGACACCATCTTCAGTAATTTC |
| seq7 | chr8:9,183,246-9,183,270 | GTCCATGTGTTAGCCACGGACATTG |
| seq8 | chr8:9,184,961-9,184,984 | GGTCTTTCCTCTATGCCAGGATTG |
| seq9 | chr8:9,183,692-9,183,715 | AACCTCCATGCAGTCGGGATGAAG |
| seqR10 | chr8:9,185,196-9,185,218 | GAATTCAAATTACAACAGACTTG |
| seqR11 | chr8:9,184,481-9,184,503 | CGCATACCCAAACGGAGTGCCAG |
| LOC promoter F1 | chr8:9,182,084-9,182,104 | CCCAGGCTTAGAGTCTATAAG |
| LOC promoter R1 | chr8:9,182,455-9,182,479 | TTCTCTTGGGATCCCCTCGGCCTTG |
| LOC157273 upstream_F1 | chr8:9,181,096-9,181,116 | GTGCCTGGCCATAGTCATCTC |
| LOC157273 upstream_R1 | chr8:9,181,748-9,181,768 | ATGTTTTCCCCCGGTTCAGAG |

* The ten primers at the top half of the table (seq2, seq3, seq4, seq5, seq6, seq7, seq8, seq9, seqR10, seqR11) target the gene-body 2.9 kb amplicon created using the long PCR primers gLOC157273_outerF2 (AGCAAAAACTGATCCGGAAG) and gLOC157273_outerR2 (TTCAGCCTCCATGGTAAAGG)

**The four primers below the line at the bottom half of the table were used to sequence the upstream 2.73 kb amplicon that was created using the long PCR primers LOCpromoterF1 (CCCAGGCTTAGAGTCTATAAG) and gLOC157273_promR1 GCTGAAATGGCTCGCTGAAC
